# BOOST: a robust ten-fold expansion method on hour-scale

**DOI:** 10.1101/2024.07.11.603043

**Authors:** Jinyu Guo, Hui Yang, Chixiang Lu, Di Cui, Murong Zhao, Cun Li, Weihua Chen, Qian Yang, Zhijie Li, Mingkun Chen, Shanchao Zhao, Jie Zhou, Jiaye He, Haibo Jiang

**Affiliations:** Department of Chemistry, The University of Hong Kong, Pok Fu Lam, Hong Kong, China; Department of Microbiology, School of Clinical Medicine, Li Ka Shing Faculty of Medicine, The University of Hong Kong, Pok Fu Lam, Hong Kong, China; Department of Geriatric Medicine and Shenzhen Clinical Research Centre for Geriatrics, Shenzhen People’s Hospital (The Second Clinical Medical College, Jinan University; The First Affiliated Hospital, Southern University of Science and Technology), Shenzhen, China; Department of Urology, The Third Affiliated Hospital, Southern Medical University, Guangzhou, China; The Third Clinical College of Southern Medical University, Guangzhou, China; Institute of Scientific Instrumentation, Shenzhen Institute of Advanced Technology, Chinese Academy of Sciences, Shenzhen, China; National Innovation Center for Advanced Medical Devices, Shenzhen, China

**Author notes:** Correspondence: Haibo Jiang.

## Abstract

Expansion microscopy (ExM) enhances the microscopy resolution by physically expanding biological specimens and improves the visualization of structural and molecular details. Numerous ExM techniques and labeling methods have been developed and refined over the past decade to cater to specific research needs. Nonetheless, a shared limitation among current protocols is the extensive time required for sample processing, particularly for challenging-to-expand biological specimens (*e.g.*, formalin-fixed paraffin-embedded (FFPE) sections and large three-dimensional specimens). Here, we have developed a rapid and robust ExM workflow named BOOST, which leverages a series of novel microwave (MW)-accelerated ExM chemistry, resulting in a single-step linear expansion of ∼10×. Specifically, BOOST facilitates a ∼10-fold expansion of cultured cells, tissue sections, and even the challenging-to-expand FFPE sections under merely 90 minutes with heat and surfactant-based protein denaturation. Furthermore, BOOST employs microwave-assisted proteomic staining and immunostaining to facilitate high-resolution visualization of structural and molecular details with significantly enhanced throughput. Noteworthily, BOOST has pioneered a ∼10-fold expansion of large millimeter-sized three-dimensional specimens in approximately three hours. BOOST offers an easily adaptable workflow based on stable and common reagents, thus boosting the potential adoption of ExM methods in biological investigations.

## Introduction

The need for better imaging methods is nearly universal for biological research. As the cornerstone of understanding biological processes, microscopy techniques have played a pivotal role in numerous scientific breakthroughs and discoveries. A recent addition is expansion microscopy (ExM) introduced by Boyden’s lab in 2015,^1^ enabling high-resolution imaging of biological samples by physically expanding the specimens, thereby overcoming the diffraction-limited resolution barrier of conventional light microscopy. Over the decade, a myriad of ExM techniques have been developed to cater to specific research aims. Recent advancements included achieving ∼25 nm resolution with a single linear expansion^2–4^ and revelation of morphological details utilizing DNA and proteomic labelling dyes.^5, 6^ When coupled with fluorescence fluctuation analysis, ExM revealed ultrastructural details of biological samples with higher resolution,^3, 7^ and even achieved the visualization of single proteins with an astonishing 1 nm resolution.^8^ The utilization of ExM techniques also revealed previously unknown assembly and organization of biological nanostructures.^9, 10^

Nevertheless, the integration of ExM on a wider scale has been constrained by the time-consuming nature of the sample preparation process. This is particularly true for challenging-to-expand biological specimens such as formalin-fixed paraffin-embedded (FFPE) sections, with the latest protease-free protocol taking around three days to expand the FFPE samples into tenfold for imaging,^3^ which preclude potential time-sensitive applications such as nanoscale pathology requiring same day diagnosis. In addition, despite progress achieved in expanding large biological samples up to fourfold their original size over extended periods (often spanning days or weeks),^11, 12^ the absence of protocols enabling higher fold expansion for large specimens remains a critical limitation, rendering unlocking hidden dimensions within large biological specimens an elusive goal. Therefore, in order to facilitate the widespread implementation of ExM, the development of a method that is rapid, robust, and compatible with a wide range of specimens is of paramount importance.

In the context of developing microscopic methods, microwave (MW) irradiation has emerged as a valuable technique for expediting sample processing. This approach speeds up both electron microscopy and optical microscopy sample preparation.^13–18^ Notably, Mayers pioneered the use of microwave in histopathology during the late 1970s, successfully fixing tissues.^19^ Subsequently, researchers explored microwave irradiation for various biological sample processing procedures, including fixation, histochemical staining, and immunohistology.^18^ Particularly, microwaves have gained recognition for antigen retrieval in paraffin-embedded tissue sections.^18^ Brief exposure to microwave irradiation considerably expedited these procedures, yielding more consistent results with enhanced histochemical and immunohistological staining.^13^ The underlying mechanisms of microwave action have been suggested to involve both generated heat and primary molecular kinetics, which contribute to the acceleration of chemical diffusion and subsequent reactions.^20^

We reasoned that the primary processes of ExM, *i.e.*, monomer infiltration, gelation, anchoring, and denaturation, could be compatible with microwave irradiation if properly designed chemistry is used as shown in other sample processing methods.^18, 21^ Also, gelation of the hydrogel monomers (*e.g.*, acrylamide) is essentially a free radical reaction known to be favored by microwave irradiation.^22^ Hence, it was deemed prudent to explore the potential of integrating microwave irradiation and developing microwave-compatible chemistry to expedite the traditionally time-consuming ExM.

Here, we developed a rapid and robust expansion microscopy workflow, BOOST, which aims to boost capability and broaden implementation in various applications. In brief, we developed microwave-compatible ExM chemistry, including anchoring chemistry, gel formulation, denaturation solutions, and labeling, together with optimized microwave irradiation protocols to expedite entire ExM workflows from gelation to final expansion. BOOST substantially reduces the processing time from several days^2–4, 23^ to less than 2 hours to 4.5 hours, depending on the sample type and staining required. In addition to the rapid processing facilitated by microwave-compatible chemistry, we demonstrated the robustness of the BOOST technique. Notably, BOOST exhibited improved gel uniformity and mechanical properties, achieved through a simplified gel formulation that eliminated sodium acrylate (SA), and avoided protease-based digestion for better preservation of epitopes for high-quality post-expansion staining.^24^ Our experiments successfully demonstrated expansion and imaging across a diverse range of biological samples, spanning sizes from a few micrometers to millimeters. These samples included cultured cells, tissue sections, FFPE sections, organoids, and whole lymph nodes, achieving an approximate tenfold expansion.

## Results

### BOOST chemistry and workflow

We developed a series of microwave-assisted chemistry and protocols for the whole workflow of ExM, from anchoring, gelation, denaturation, wash and expansion. The running time for various sample types varies while shortening the expansion process for all sample types to under two hours (**Fig. 1A**). When compared to other protocols to date ^2–4, 23, 25^, there has been a significant reduction in the amount of time used, making it about 6-40 times faster (**Fig. 1B**), while achieving ∼10-fold single linear expansion factor. Hence, we termed this newly developed ExM protocol as BOOST.

**Fig. 1:**
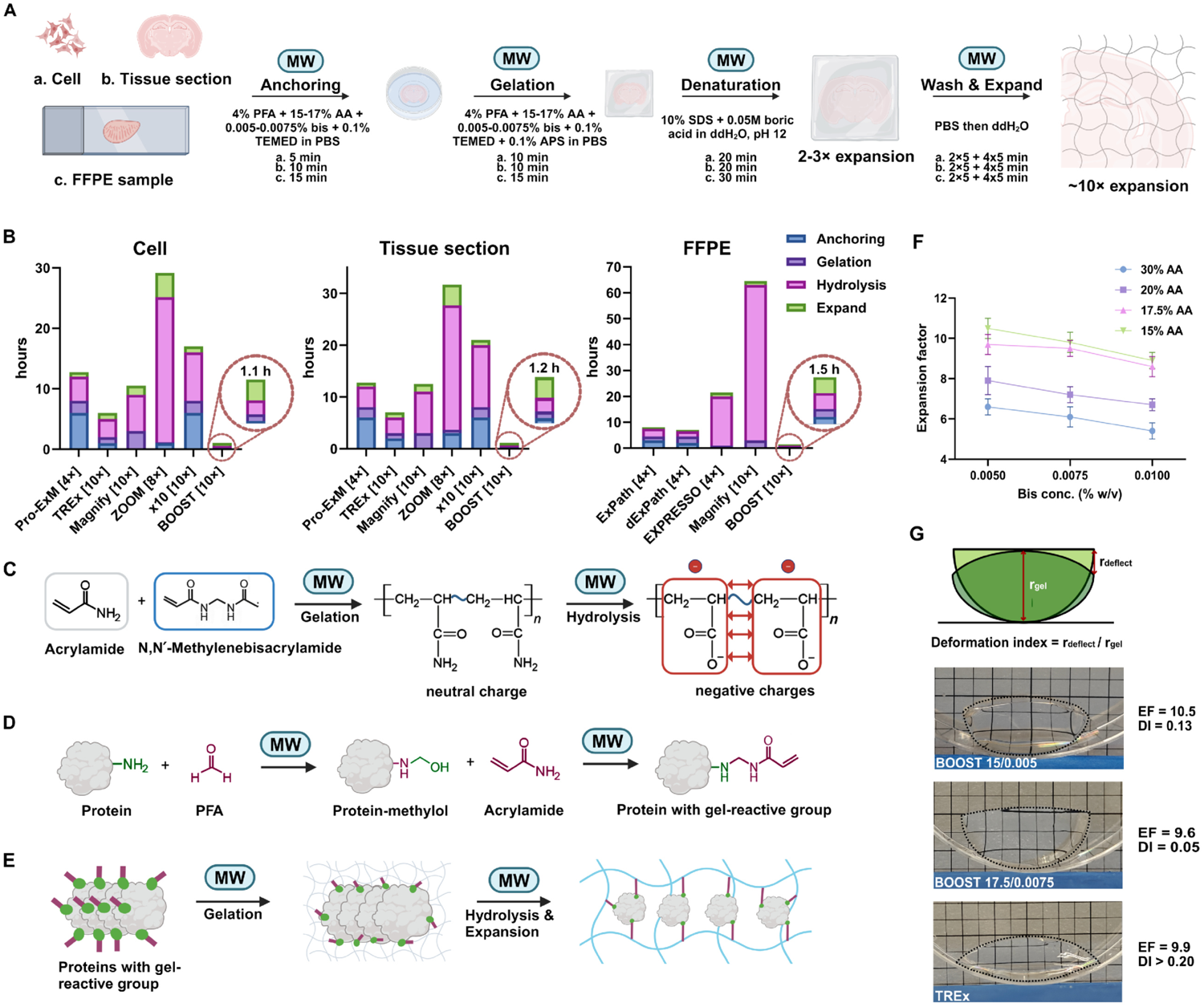
Development of BOOST workflow. **(A)** Optimized BOOST microwave-assisted protocols for three sample types. Reagents and processing times are listed for each step. **(B)** Comparison of processing times for selected ExM techniques for three types of samples. **(C)** Microwave-assisted BOOST hydrogel chemistry uses free-radical initiated polymerization of AA and bis, followed by alkaline hydrolysis to convert amide groups into charged carboxyl groups, transforming the gel into a swellable polyelectrolyte hydrogel. **(D)** Microwave-assisted anchoring with the reactions between the amine groups of the biomolecules and aldehyde of PFA to produce methylol groups, followed by reactions with AA to be anchored to the hydrogel. **(E)** A schematic diagram of biomolecules undergoing microwave-assisted gelation, hydrolysis, denaturation, and expansion; green dots and purple sticks depict the anchorable gel-reactive groups on biomolecules. **(F)** Optimization of concentrations of AA and bis for maximizing the expansion factor (EF) under 20-minute microwave-assisted hydrolysis (mean ± SD, n = 3). **(G)** Gel mechanical performance measurement showing the deformation indexes of BOOST gel formulations and the TREx gel. Squares in the background indicate 5 mm × 5 mm areas.

At the early stage of the development, we selected the hydrogel chemistry without SA addition due to many reported problems.^4, 25, 26^ An acrylamide-only gel recipe introduced by ZOOM^25^ based on 30% (w/v) acrylamide (AA) and a cross-linking agent 0.01% (w/v) N,N’-methylenebisacrylamide (bis) was initially employed here to develop the early microwave-compatible gelation and denaturation processes. For the gelation process, like conventional polyacrylamide gels, bis crosslinks polymer chains to the network under free radical polymerization,^27^ which can be accelerated by low-energy microwave irradiation^22^ and we subsequently confirmed this by carrying out the gelation process under microwave using the acrylamide-only gel recipe (**Fig. 1C**). For the denaturation process, as SA is absent in the crosslinked hydrogel, simultaneous hydrogel hydrolysis is required to transform the neutral polyacrylamide gel into a swellable polyelectrolyte hydrogel, facilitating the expansion process (**Fig. 1C**), which can also be theoretically assisted with microwave irradiation under alkaline conditions.^28^ To validate, we tested hydrolysis of the crosslinked blank hydrogel (30% w/v AA + 0.01% w/v bis) with an alkaline buffer with pH ranging from 9 to 12 under microwave irradiation, a maximal pH of 12 was chosen here for 12 is the highest pH reported for non-destructive treatment proteins.^29^ We observed an increase in expansion factor (EF) with the increasing pH, reaching a maximum of ∼5× EF (Fig. S1A). Our findings also indicate that a 20-minute duration is sufficient to achieve the maximum EF for this specific gel formulation (Fig. S1B). Simultaneous to the hydrogel hydrolysis, anchored proteins need to be digested to ensure uniform expansion. We optimized a denaturation method based on heat and chaotropic reagent, sodium dodecyl sulfate (SDS), which is a recognized method from SDS-PAGE^30^ and microwave compatible,^21^ and is being suggested to facilitate better epitope preservation for better immunostaining.^26, 31^ We therefore also employed 10% w/v SDS in the hydrolysis buffer. Toward intact morphology and protein preservation, we then evaluated the impact of hydrolysis buffer pH on the preservation of morphology and proteins with samples embedded. We performed N-Hydroxysuccinimide (NHS) ester staining on ZOOM-processed mouse liver sections and observed no significant differences in protein intensity or morphology across the tested pH range under microwave hydrolysis (pH 10 to pH 12) (Fig. S1C).

In addition to the microwave-adapted gelation and hydrolysis step, we set out to seek a microwave-compatible anchoring strategy to maximize biomolecular tethering to the hydrogel within a limited time. To start, we questioned whether the widely used anchoring strategy by Acroylol-X (AcX)^26^ would be compatible with microwave irradiation. The Ten-fold Robust Expansion Microscopy (TREx) protocol^2^ was used in this case with only the anchoring step being modified into several testing conditions (Fig. S2A). Notably, microwaved samples exhibited inferior preservation compared to the standard AcX-incubated samples. Instead, the microwaved samples resembled the one without any anchoring, suggesting that the acceleration of AcX and amine reactions was ineffective under microwave conditions (Fig. S2A). Next, we tested different anchoring strategies (Fig. S2B and finalized with a microwave-assisted anchoring strategy with AA + PFA (**Fig. 1D**). In this anchoring strategy, amine groups on biomolecules are initially modified by PFA to form a reactive methylol group, which further reacts with surrounding excess acrylamide monomers to form a protein with a gel-reactive group that is later anchored to the gel matrix during polymerization (**Fig. 1E**).^24, 32^ The acceleration of anchoring methods using PFA and AA is consistent with previous uses of PFA for the protein fixation with improved effectiveness and uniformity under microwave irradiation^33^, ensuring that reactive methylols are rapidly available for further acrylamide reactions.

To achieve higher resolution, we further developed the gel formulation to increase EFs while preserving adequate gel mechanical properties. We tested gel formulations with various AA and bis concentrations (**Fig. 1F**). The mechanical property (measured by deformation index (DI) and corresponding expansion factor of blank gels of the selected formulas were measured and calculated^2^ (**Fig. 1G**), and the lowest DI of 0.05 was achieved by 17.5% w/v AA/0.0075% w/v bis recipe with an EF of 9.6×, and the largest EF of 10.5× was achieved by 15% w/v AA/0.005% w/v bis formula with DI of 0.13. Both DIs indicate an overall stronger gel compared with the TREx formula, which obtained a DI of above 0.20 at a 10× expanded state (**Fig. 1G**). The results obtained from the DI values of the selected formulas (0.05 and 0.13) demonstrate that they exhibit satisfactory mechanical strength.^2^ The highest EF formula is potentially ideal for cultured cell samples, whereas the high mechanical strength formula could be used for larger tissue samples, which would need firmer gel for handling and transfer.

### BOOST reveals ultrastructural details of cultured cells, tissue sections, and FFPE samples

To visualize the overall high-quality morphology with less time spent on staining, we optimized microwave-compatible proteomic covalent staining with NHS esters and nucleic acid staining with SYTOX Green for a wide range of biological samples (*e.g.*, cultured cells, tissue sections, and mouse and human FFPE samples). We achieved 4-12 times faster morphological staining than reported methods (**Fig. 2B**). ^2, 3, 5, 34^ ^6^ More importantly, we improved the uniformity of morphological staining across thicker samples, likely due to the microwave-compatible crosslinking reactions between NHS esters and primary amines^35^ and increased diffusion enabled by microwaves.

**Fig. 2:**
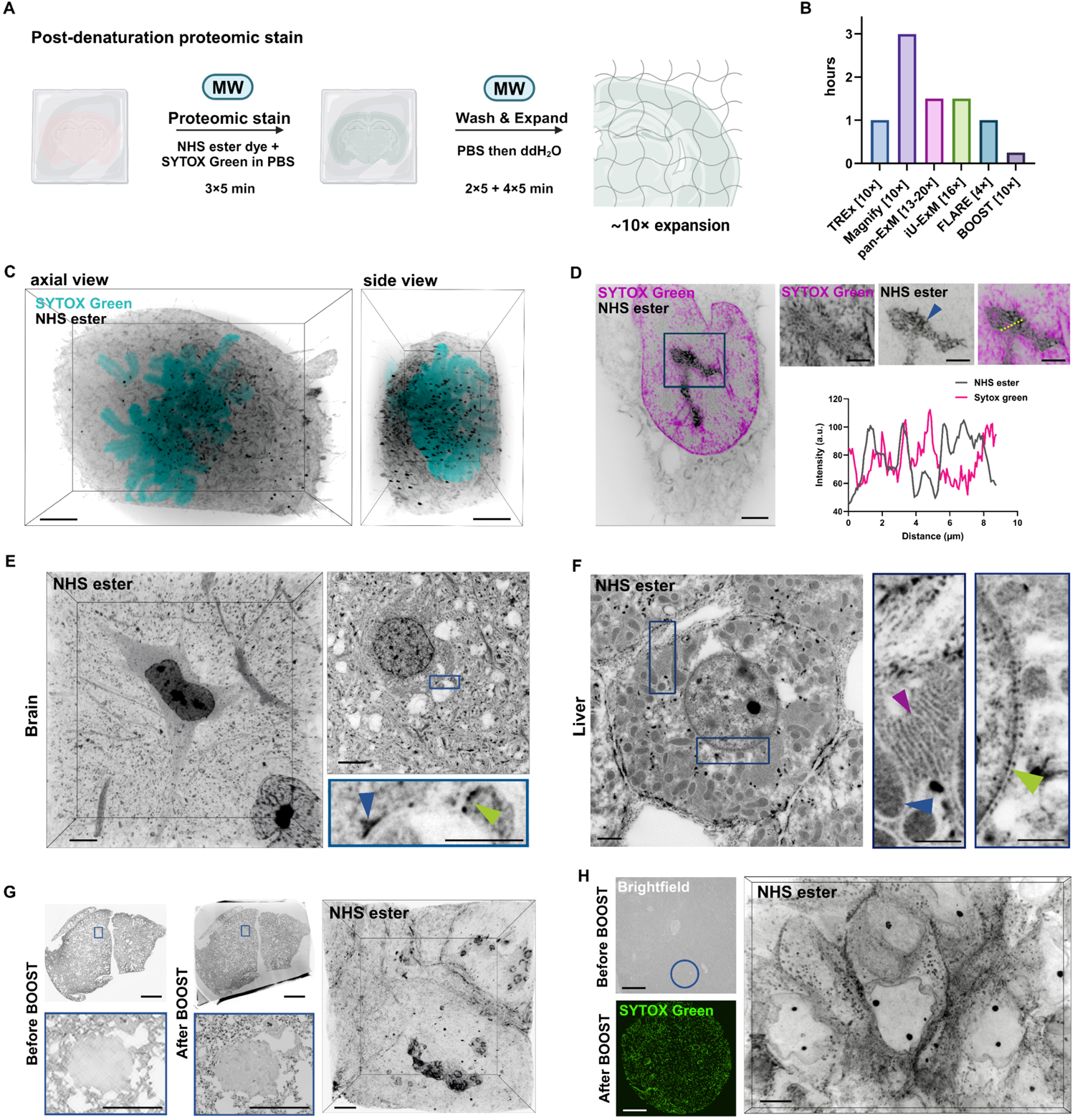
Ultrastructural staining on BOOST processed samples. **(A)** Schematic of ultrastructural staining after BOOST denaturation with reagents and processing times listed. **(B)** Comparison of the BOOST microwave-assisted NHS ester staining time with ExM techniques that used post-processing NHS ester staining. **(C)** 3D visualization of a mitotic U2-OS cell stained with ATTO 565 NHS ester and SYTOX Green, EF = 10.2×. **(D)** *Left*, 2D image of a U2-OS cell stained with ATTO 565 NHS ester and SYTOX Green, EF = 10.2×. *Right*, magnified views of the boxed region. *Blue arrow*, granular component of nucleolus. Line profile along the dashed line indicating a partially complementary staining pattern of NHS ester and SYTOX Green in the nucleolus. **(E)** 3D and 2D view of neurons in a mouse brain section stained with ATTO 488 NHS ester, EF = 9.1×. A magnified view of the boxed region shows synaptic active zone (*blue arrow*) and synaptic vesicles (*green arrow*). **(F)** 2D image of expanded mouse liver section stained with ATTO 488 NHS ester, EF = 9.1×. Two magnified views of the boxed regions showing endoplasmic reticulum (*purple arrow*), mitochondria (*blue arrow*), and nuclear pores (*green arrow*). **(G)** Images of an ATTO 488 NHS ester stained FFPE section of mouse lung bearing tumors before and after BOOST treatment. Post-BOOST images were taken by shrinking the expanded section (8.9×) back to ∼2.6× in PBS. Magnified views of the boxed region showing a tumor before and after expansion. *Right*, a 3D visualization of the expanded mouse tumor, EF = 8.9×. **(H)** Images of a human prostate cancer FFPE section before and after BOOST. A pre-BOOST processing image was taken using brightfield microscopy. Post-BOOST image of an expanded punched-out circular region (4 mm), and the section was stained with SYTOX Green, EF = 8.6×. *Right*, 3D visualization of the expanded tumor region stained with ATTO 565 NHS ester. Scale bars (biological scale): 2 μm (C); 2 μm (D, *left*); 1 μm (D, *right*); 2 μm (E, *left* and *top right*); 1 μm (E, *bottom right*); 2 μm (F, *left*); 1 μm (F, *middle* and *right*); 1 mm (G, *top left* and *top middle*); 400 μm (G, *bottom left* and *bottom middle*); 2 μm (G, *right*); 100 μm (H, *top left*); 200 μm (H, *bottom left*); 2 μm (H, *right*).

For cultured cells, we stained U2-OS cells with ATTO 488 NHS ester under two staining conditions (*i.e.*, 1 hour at room temperature as previous protocols suggested^2, 3, 5^ and 10 minutes under microwave irradiation). We found no significant differences in the staining (Fig. S3A), both revealed ultrastructural details of organelles such as mitochondria by their morphological characteristics. We then pondered upon the choice of NHS ester dyes with different hydrophobicity and potential differential compartment labeling.^36^ We tested three NHS esters that are conjugated with ATTO dyes (ATTO 488, ATTO 565, ATTO 647N) owning different hydrophobicity on expanded U2-OS cells. We observed variations in the intensity of the nucleus labeling amongst the three, but all NHS esters were compatible with microwave irradiation and showed no major difference in the mitochondria structure visualization (Fig. S3B). We then established the post-BOOST proteomic staining by NHS ester dyes with simultaneous SYTOX Green staining under the microwave. The staining time is 15 minutes for all three types of samples (**Fig. 2A**), which is considerably shorter than other protocols for post-processing proteomic staining (**Fig. 2B**). Using BOOST with the proteomic staining strategy, we visualized a mitosis U2-OS cell with down to 25 nm resolution (**Fig. 2C and supplementary video 1**). We also visualized that SYTOX Green and NHS ester staining showed complementary structures of nucleolus, outlined by the differential compartment labeling in the nucleolar granular components (**Fig. 2D**).

For tissue sections, we demonstrated rapid and homogenous NHS ester staining on the mouse brain, kidney, and liver 100-µm sections and revealed ultrastructural information in thick (100 µm) sections (**Fig. 2E-2F**, Fig. S4A). We imaged a mouse brain section and a 3D structure of a neuron in the cortex was revealed with down to ∼30 nm resolution (**Fig. 2E and supplementary video 2**). The boxed region showed a zoomed-in view of a brain synapse with heavy dots representing resolved synaptic vesicles and a heavily curved stained active zone (**Fig. 2E**). In mouse liver sections stained with ATTO 488 NHS ester (**Fig. 2F**), the nucleus of each hepatocyte was outlined with high labeling in the nucleoli. The nuclear envelope showed dense labeling on the inner nuclear membrane with heavily stained nuclear pore complexes (**Fig. 2F**). Several organelles in the cytosol, such as endoplasmic reticulum (ER) and mitochondria, were clearly visible (**Fig. 2F**). Unlike the high-quality lateral visualization, the uniform staining of thick tissue throughout the thickness always poses an issue, even staining at the expanded state.^6^ Notably, we observed a greatly increased staining uniformity of a 100-μm thick section under 15-minute microwave processing compared to 1-hour incubation under standard conditions (Fig. S4B), likely due to the microwave enables increased diffusion and simultaneously, the NHS ester and amine reaction rate.^35^ In addition to the single NHS strategy, the dual NHS ester labeling approach was found to label complementary ultrastructures^36^ and potentially enable better visualization of morphological contrasts. We tested dual NHS ester labeling for expanded tissue sections such as brain (Fig. S5A) and kidney (Fig. S5B). We observed different NHS ester stained different structures (Fig. S5), in particular, ATTO 565 NHS ester showed strong labeling signals on the nucleus (Fig. S5).

Importantly, we optimized BOOST for FFPE samples. As FFPE samples have a highly cross-linked nature,^37^ we integrated a step of 30-minute microwave-assisted antigen retrieval step in Tris-EDTA buffer (pH 9) before anchoring to reverse heavy chemical crosslinks. The role of microwave treatment for antigen retrieval has been established.^15, 38^ We verified the importance of the antigen retrieval step for the BOOST; absence or insufficient antigen retrieval would lead to insufficient anchoring, therefore, fractures at an expanded state (Fig. S6). We tested the mouse lung tumor FFPE sample and found the whole section was well expanded by BOOST, with the ultrastructural details of the tumor region being clearly discerned (**Fig. 2G**). We then tested BOOST on the human prostate tumor FFPE samples; after homogeneous expansion of the sample to an 8.9× fold, the overall 2D and 3D morphology of the tumor region was also revealed using NHS ester staining (**Fig. 2H**).

### BOOST is compatible and accelerates immunostaining

We optimized microwave-accelerated immunostaining for three types of gel-embedded biological samples (*i.e.*, cultured cells, tissue sections, and FFPE sections) (**Fig. 3A**). Microwave-accelerated immunohistostaining, immunocytochemistry, and immunofluorescence have long been tested^14, 17, 39,40^ and benefits with microwave irradiation during antigen-antibody incubation were also verified.^16^ We modified the previously reported microwave immunostaining protocol^40^ and achieved the microwave-assisted immunostaining of gel-embedded samples within an hour (**Fig. 3B**), which was ∼10-80 times faster than other protocols for immunostaining of gel-embedded samples.^3, 23, 25, 34^

**Fig. 3:**
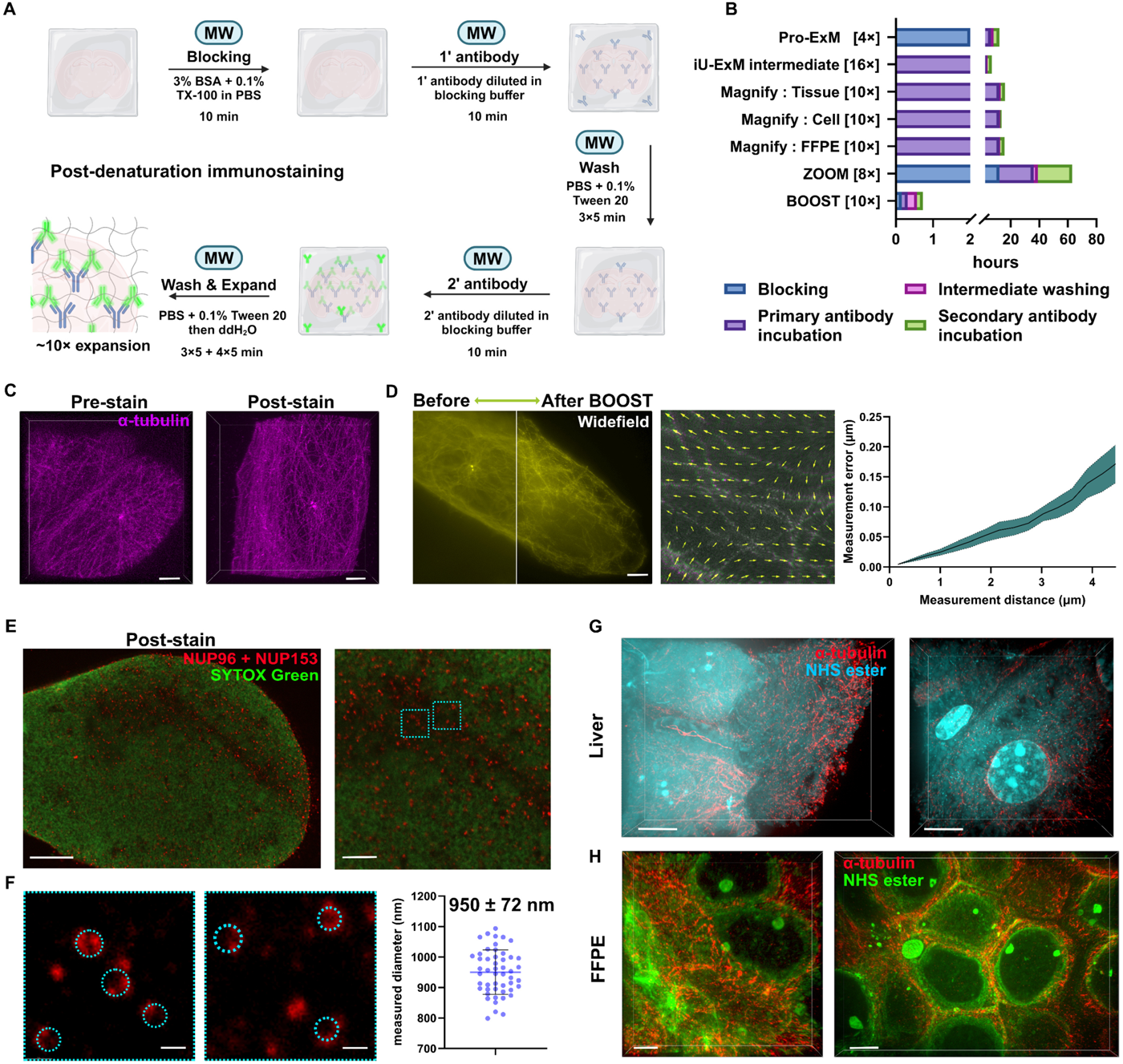
BOOST is compatible with rapid immunostaining on various sample types. **(A)** Schematic of microwave-assisted immunostaining after BOOST denaturation with the reagents and processing times listed. **(B)** Comparison of the immunostaining duration with selected ExM techniques that used post-denaturation immunostaining. **(C)** Images of expanded U2-OS cells stained for α-tubulin before BOOST (EF = 9.5×) or after BOOST (EF = 9.1×). Immunostaining after BOOST denaturation showed a significantly improved signal-to-noise ratio. **(D)** Distortion measurements with epifluorescence images of U2-OS cells stained for α-tubulin before and after the BOOST processing, EF = 9.5×. *Middle*, A distortion field with RMS error plot of α-tubulin images before and after BOOST processing. *Right*, the estimated distortion error of ∼4% of the measured length (n = 6) indicating minor distortions. **(E)** Post-BOOST immunostaining of NPCs of a U2-OS cell with anti-NUP153 and anti-NUP96 antibodies, and DNA was stained with SYTOX Green, EF = 9.6×. **(F)** Magnified views of the boxed regions showing the structures of individual NPC. The dashed circles were manually fitted on the NPC staining and quantified to plot the mean diameter of NPCs with standard deviations (n = 50 NPCs from three distant cells). **(G)** 3D views of an expanded mouse liver section stained with α-tubulin and ATTO 488 NHS ester, EF = 8.9× **(H)** 3D views of expanded human prostate cancer FFPE section stained with α-tubulin and ATTO 488 NHS ester, EF = 8.7×. Scale bar (biological scale): 2 μm (C); 2 μm (D); 2 μm (E, *left*); 500 nm (E, *right*); 100 nm (F); 2 μm (G, *left*); 1 μm (G, *left*); 1 μm (H, *left*); 2 μm (H, *right*).

We verified that BOOST is compatible with microwave-assisted immunostaining both before and after the expansion procedure, while immunostaining after the expansion procedure achieved improved staining results with clearly resolved microtubule structures (**Fig. 3C and supplementary video 3**). This is consistent with works of molecular decrowding with ExM to improve epitope accessibility and to get better immunostaining results.^9^ To ensure the structures were uniformly expanded, we quantified the error introduced by expansion by comparing microtubules immunostained before and after BOOST with the same microscope setup. The distortion measured was under 5% of the measurement length, indicating low distortions (**Fig. 3D**). We reasoned immunostaining after BOOST would be a preferred method to achieve improved staining though immunostaining before BOOST was also feasible.

We also validated BOOST by staining other structures in cells that were not observable using confocal microscopes. We demonstrated the visualization of nuclear pores using SYTOX Green or DAPI staining and immunostaining with antibodies for nuclear pore complex (NPC) (anti-NUP 153 and anti-NUP 96) (**Fig. 3E**, Fig. S7A-B). We used a gel formula of 17.5% AA and 0.0075% bis and achieved an EF of 9.6× determined with the macroscopic method. We quantified 50 NPCs chosen from three distant cells (**Fig. 3F**) and obtained NPC sizes of 950 nm ± 72 nm (99 nm ± 8 nm when translated into original size using EF = 9.6). This further translates the local EF of 8.9× calculated from NPC sizes of 107 nm, with a 7.4% deviation from the EF determined with the macroscopic method, which is more than acceptable as suggested by TREx.^2^

Next, we verified the immunostaining procedure for other types of samples. We combined NHS ester staining with immunostaining against α-tubulin on the BOOST-processed mouse liver sections and clearly observed the organization of microtubules within the liver tissues (**Fig. 3G**). Outstandingly, BOOST also achieved rapid expansion and staining on human prostate cancer FFPE sections (**Fig. 3G**). We further validated the microwave-accelerated staining results for the human pathology samples, as the accuracy will be critical for pathological investigations. We observed identical staining patterns with an α-tubulin antibody and a prostate-specific membrane antigen (PSMA) antibody, comparing with microwave-accelerated staining and conventional protocols performed for two days (Fig. S7C).

### BOOST is capable of rapidly expanding large samples to ∼10-fold

We further adjusted the anchoring, gelation, hydrolysis, and wash and expand steps of BOOST to expand larger biological samples with original sizes ranging from hundreds of micrometers to a few millimeters and achieved ∼10× expansion and obtained millimeter-to-centimeter-size expanded samples, which has not been reported by any current protocols. Here, we demonstrated the homogenous expansion of organoids and whole lymph nodes, which only took ∼2 hours and ∼3 hours, respectively (**Fig. 4A**).

**Fig. 4:**
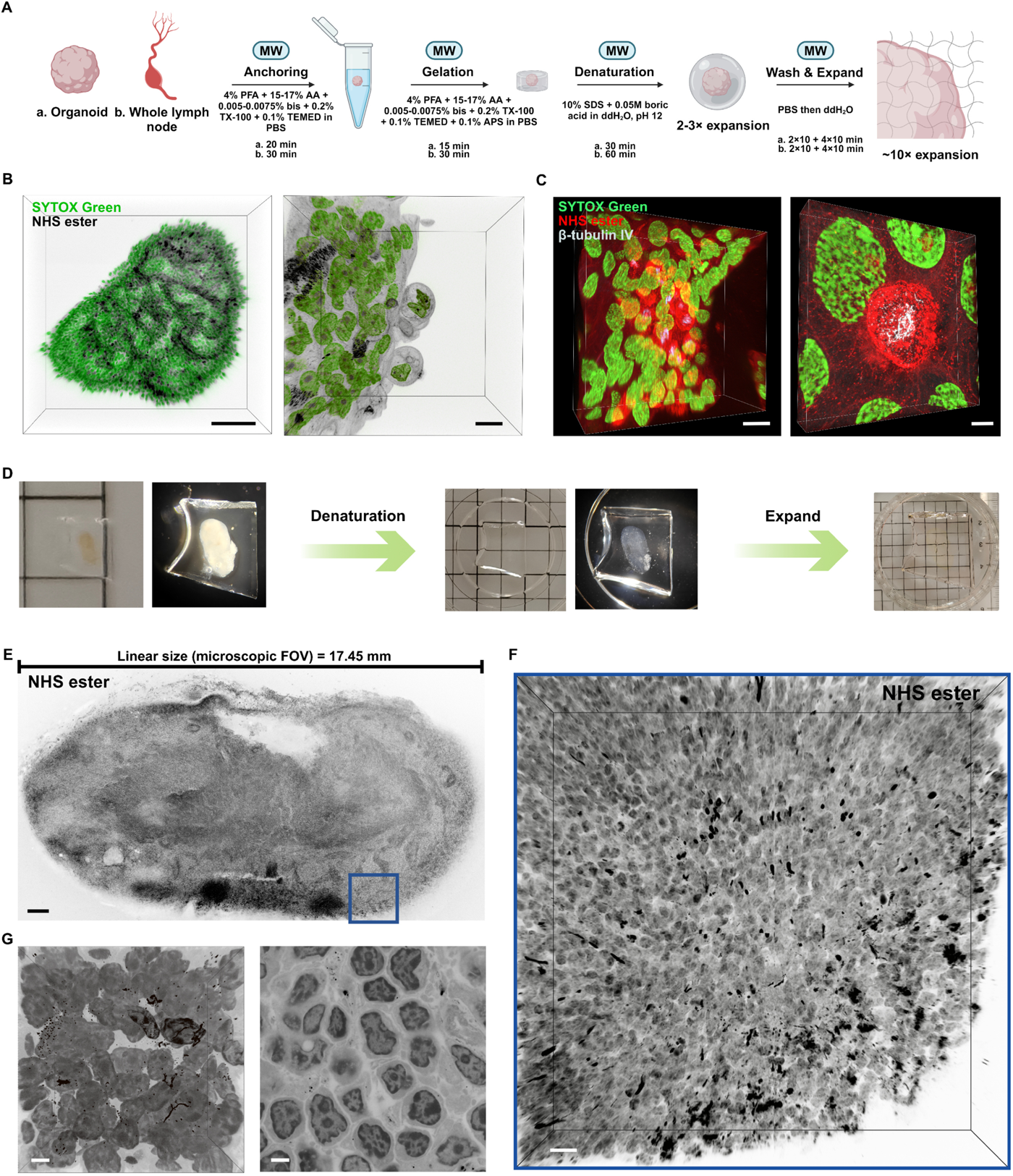
BOOST rapidly expands large biological specimens. **(A)** Schematic of BOOST workflow for large biological specimens with reagents and processing times listed. **(B)** 3D visualization of an expanded whole airway organoid stained with ATTO 565 NHS ester and SYTOX Green, and a magnified view of the airway organoid periphery, EF = 10.1×. **(C)** 3D visualization of a selected region of the airway organoid stained with ATTO 565 NHS ester and SYTOX Green, and immunostained for β-tubulin IV. **(D)** Processing workflow of an intact lymph node to achieve a 9-fold expansion. Each square in the background represents a 5 mm × 5 mm area. (**E**) 2D image of the middle section of intact lymph node stained with ATTO 565 NHS ester, and (**F**) a magnified 3D view of the boxed region, EF = 9×. **(G)** 3D and 2D view of selected regions in the intact lymph node. Scale bar (biological scale): 100 μm (B, *left*); 10 μm (B, *right*); 10 μm (C, *left*); 2 μm (C, *right*); 100 μm (E); 10 μm (F); 2 μm (G).

Specifically, we demonstrated BOOST to characterize the cellular composition and organization in three dimensions and morphology and ultrastructure of ∼500-µm airway organoids with an EF of 10.1× when employing 15% w/v AA / 0.005% w/v bis gel formulation. With the staining of SYTOX Green and NHS ester, we reconstructed the 3D morphology of the whole organoid in detail (**Fig. 4B**, **supplementary video 4**, and **supplementary video 5**). Combining microwave-assisted immunostaining, we further revealed the distribution of β-tubulin IV in the lung airway organoid with ∼30 nm resolution (**Fig. 4C**).

We further demonstrated the capability of BOOST to expand even larger samples, using lymph nodes in the range of several millimeters. The robustness of BOOST enabled millimeter-sized samples to achieve an EF beyond just 4× by existing techniques involving millimeter-sized sample.^11, 12^ We adjusted the BOOST protocol in the processing time for sufficient anchoring, gelation, hydrolysis, and staining to achieve whole lymph node visualization at ∼9× fold with 17.5% w/v AA / 0.0075% w/v bis gel formulation in ∼4.5 hours (**Fig. 4A**). We demonstrated the embedding of a lymph node with a measured vertical size of 1.95 mm; the lymph node was later expanded to 2.8-fold in PBS after denaturation/hydrolysis; and a further expansion in water led to a final vertical size of ∼1.75 cm, indicating a 9× macroscopic EF with almost transparent appearance under normal lighting (**Fig. 4D**). We then combined proteomic staining to image the 3D morphology of the whole lymph node in detail; we imaged across the middle region of the lymph node to confirm the uniformity of the NHS ester labeling (**Fig. 4E**). A further magnified region was imaged to visualize morphological details of the expanded lymph node (**Fig. 4F-4G**). Hence, BOOST achieved a ∼10-fold expansion of millimeter-sized three-dimensional samples that was not feasible before.

## Discussion

We developed BOOST—a new hour-scale and robust workflow for expanding diverse biological specimens to boost ExM’s throughput and adoption in biological investigations (Fig. S8). BOOST has made significant advances in the following aspects: (1) BOOST has dramatically improved the throughput of ExM by innovatively using a series of microwave-assisted chemistries for all major ExM procedures (*i.e.*, monomer infiltration, anchoring, gelation, denaturation, staining, and expansion). (2) BOOST is highly robust and versatile. It uses common reagents and instruments, and it works effectively on various sample types, including cultured cells, tissues, organoids, and FFPE samples, and is also compatible with morphological staining and immunostaining. (3) BOOST has pioneered the single-step ∼10-fold linear expansion of millimeter-sized large 3D biological specimens.

The improved throughput of BOOST was achieved largely due to the innovative implementation of microwave-compatible ExM chemistries. Microwave irradiation accelerates mainly monomer infiltration, anchoring, gelation, denaturation/hydrolysis, and staining chemistries, with the most significant acceleration for the denaturation step. The improvement in the throughput was particularly evident for expanding FFPE samples, as previously a homogenous linear expansion of 10-fold for FFPE samples was achieved by Magnify. BOOST reduced the whole workflow from days in Magnify^3^ to less than 4 hours, inclusive of staining for ultrastructure and proteins of interest. The significant enhancement in throughput may facilitate time-sensitive applications, such as nanoscale pathology. Researchers have previously utilized ExM techniques to obtain ultrastructural details in renal pathological samples yet only with 4-fold expansion.^7, 41^ This reduction in processing times, when coupled with 10-fold expansion, could provide a crucial advantage for clinical applications.

The robustness and versatility of BOOST were demonstrated by the gel rigidity, non-proteolytic digestion, and compatibility with various samples and different staining protocols. Importantly, BOOST utilized a simple gel formulation with only AA and bis and avoided SA, a gel monomer present in most ExM gel formulations to date.^26^ The absence of SA helps to avoid potential issues, such as variable impurities from vendors, potential tissue shrinkage, and uneven distribution of ionic residues, as reported by other studies.^4, 25, 26^ The use of heat and chaotropic reagents for denaturation improved epitope preservation and enabled high-quality post-expansion staining facilitated by protein decrowding.^24, 26, 31^ All the reagents used in the BOOST workflow are reliable and easily accessible, which could potentially boost the adoption of the technique for biological investigations.

It is noteworthy that BOOST successfully accomplished a single-step ∼10-fold expansion of large 3D biological specimens, a task previously unattainable. This improvement is likely attributable to the enhanced monomer infiltration, anchoring, homogenous gelation, and denaturing within large specimens facilitated by microwave irradiation. In contrast to conventional 2D imaging of tissue sections, 3D imaging excels in capturing the intricacy of biological samples. The advanced capabilities of 3D imaging are crucial for comprehending cellular composition, and interactions between cells in intact biological samples.

In developing the current BOOST protocol, our primary focus was on enhancing the speed and reliability of major ExM steps. To foster the future adoption of BOOST in both research and clinical settings, several potential advancements could be explored. For instance, we have demonstrated compatibility with immunostaining and preservation of proteins. Validation of the preservation of other biomolecules, such as RNA, should be considered. The application of microwave-assisted processing may also be extended to other steps of the workflow, including long-verified sample fixation and washing.^18^ Additionally, the integration of computational methods with BOOST could potentially enhance resolution to a further scale, which may prove advantageous for specific applications.

## Acknowledgments

This work is supported by the Hong Kong Research Grant Council General Research Fund (17102722, 17300523), and National Natural Science Foundation of China (32271445) to H.J. We thank the Imaging and Flow Cytometry Core at the Centre for PanorOmic Sciences (CPOS) at HKU for the technical support in imaging acquisition. The work was conducted in the JC STEM Lab of Molecular Imaging, funded by The Hong Kong Jockey Club Charities Trust.

## Author contributions

HJ and JG designed the experiments and wrote the paper. All performed experiments, collected, analyzed, and assembled the results. HJ secured funding. All have commented on and edited the manuscript.

## Figures and Figure Legends

**Fig. S1:**
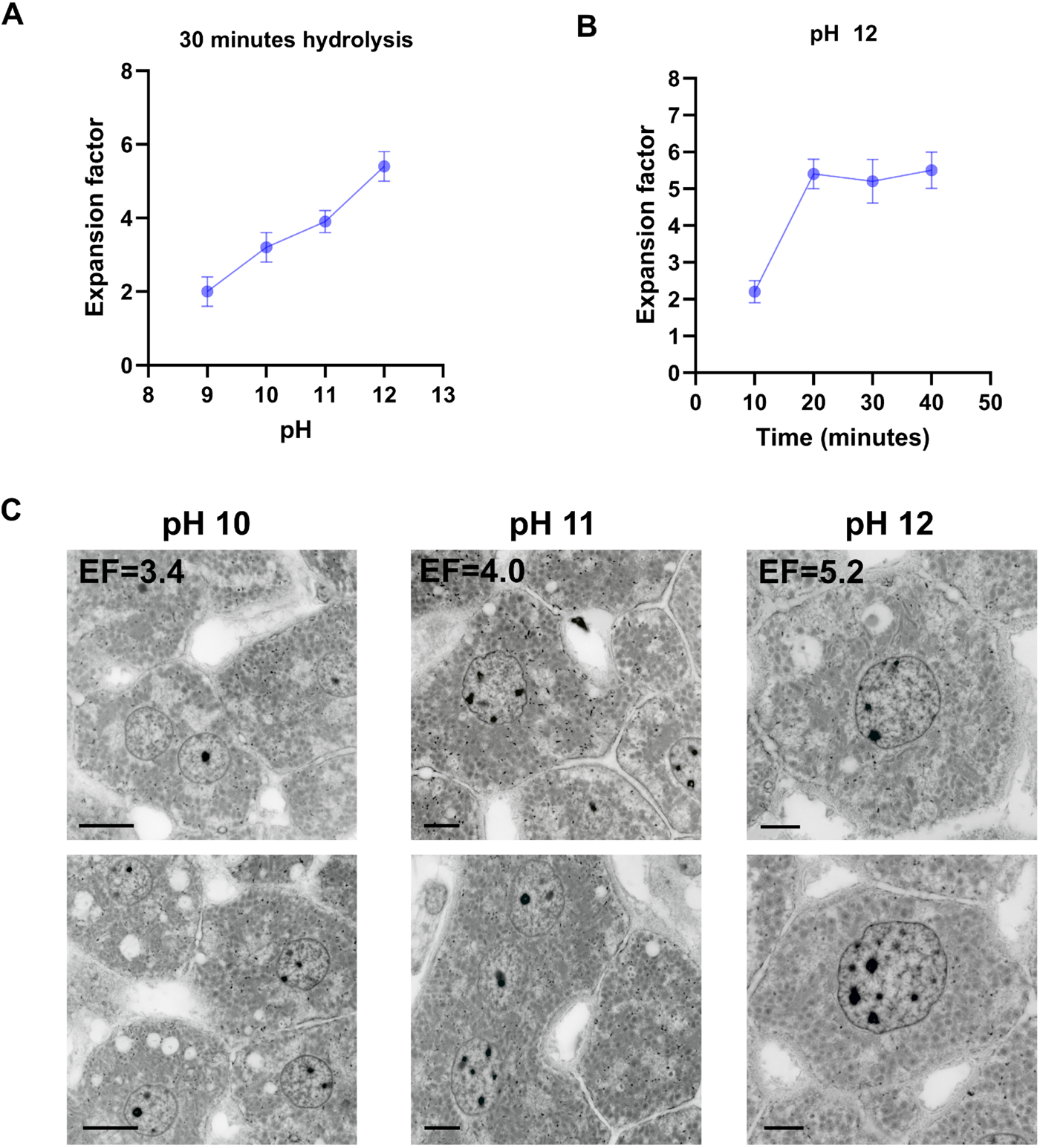
Developing BOOST hydrolysis strategy. **(A)** Optimization of the pH of the hydrolysis buffer to maximize the EF under microwave irradiation for 30 minutes with 30% w/v AA and 0.01% w/v bis blank gels (mean ± SD, n = 3). **(B)** Optimization of the microwave-assisted hydrolysis processing times to achieve the highest EF with pH 12 hydrolysis buffer (mean ± SD, n = 3). **(C)** Assessment of the morphology preservation and NHS ester labeling efficiency with 100-µm mouse liver sections under various alkaline strengths. All samples were stained with ATTO 565 NHS ester. Scale bars (biological scale): 10 µm (*left*); 5 µm (*middle*); 4 µm (*right*).

**Fig. S2:**
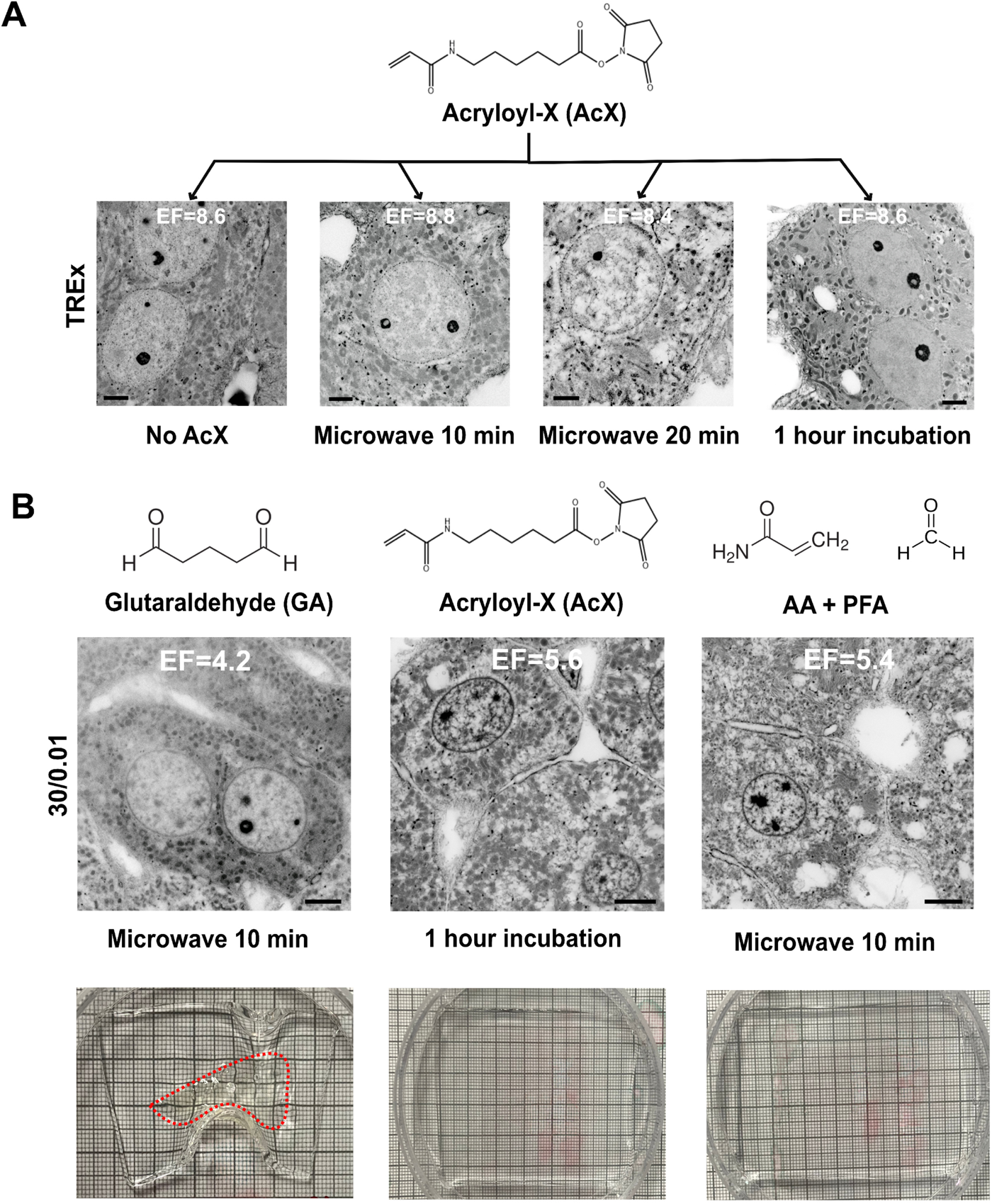
Developing BOOST anchoring strategy. **(A)** Assessment of the compatibility of AcX anchoring using mouse liver sections under microwave irradiation of 10 minutes or 20 minutes, and no anchoring and TREx’s AcX anchoring protocol was used as a control. All samples were processed with TREx protocols. Samples showed less preserved morphology under AcX anchoring with microwave irradiation. **(B)** Evaluation of anchoring strategies comparing glutaraldehyde (10-minute microwave irradiation), AcX (1-hour room temperature incubation), and AA+PFA (10-minute microwave irradiation) based on gel formula of 30% w/v AA and 0.01% w/v bis. Microwave-assisted AA+PFA anchoring showed a similar preservation of the morphology to the standard TREx AcX anchoring method. All samples were mouse liver section samples and stained with ATTO 565 NHS ester to visualize morphology preservation. Scale bars (biological scale): 2 µm (A); 5 µm (B).

**Fig. S3:**
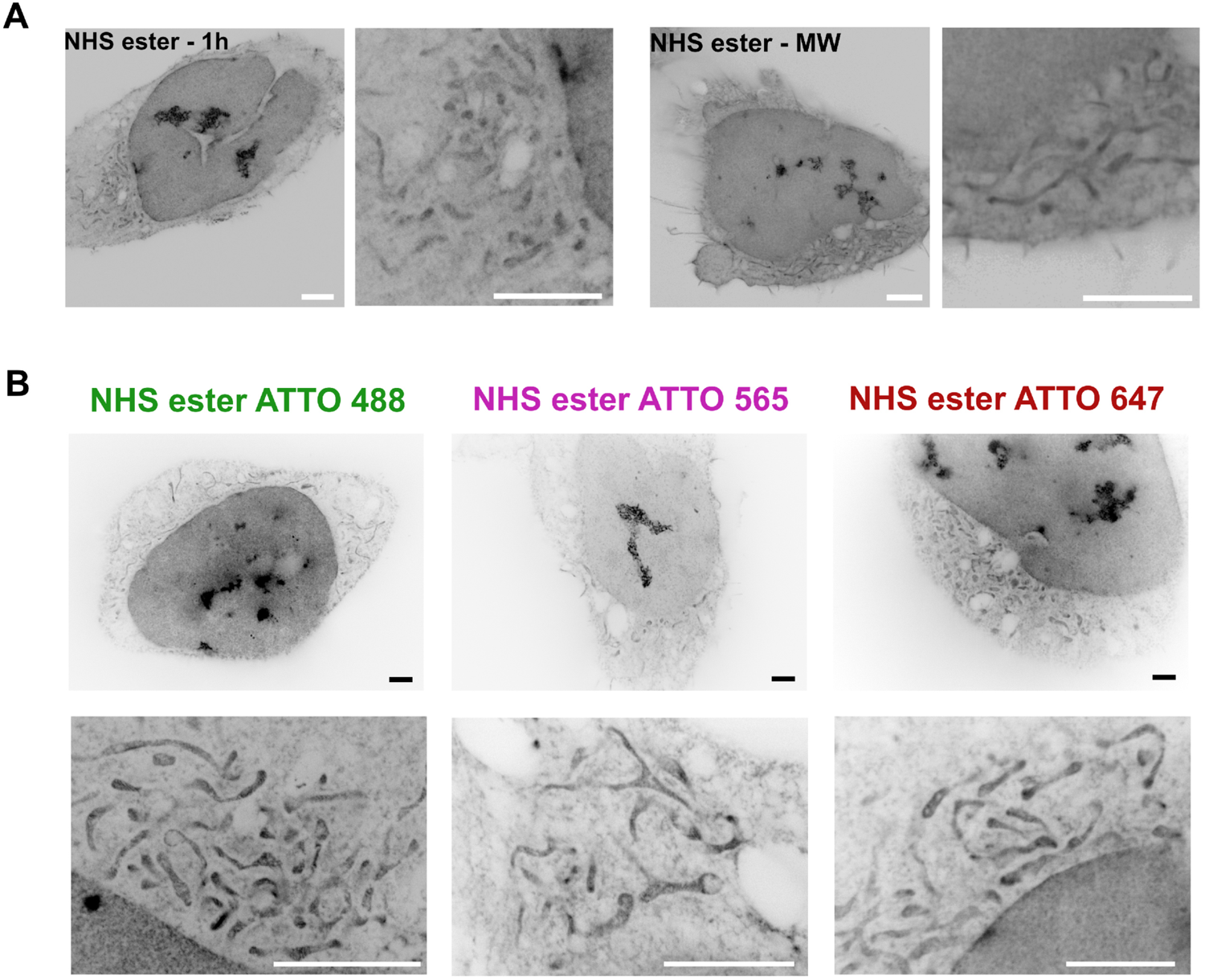
Developing BOOST morphological staining strategy for U2-OS cells. **(A)** Comparison between standard 1-hour NHS ester incubation (EF = 9×) and microwave-assisted 15-minute NHS ester incubation (EF = 9.1×), and both enabled the visualization of mitochondria in a subcellular context. **(B)** Comparison of staining with different NHS esters for microwave-assisted morphological staining. All three NHS esters enabled visualization of the mitochondria with slight differential staining of subcellular structures with the three NHS esters. Scale bars (biological scale): 2 µm.

**Fig. S4:**
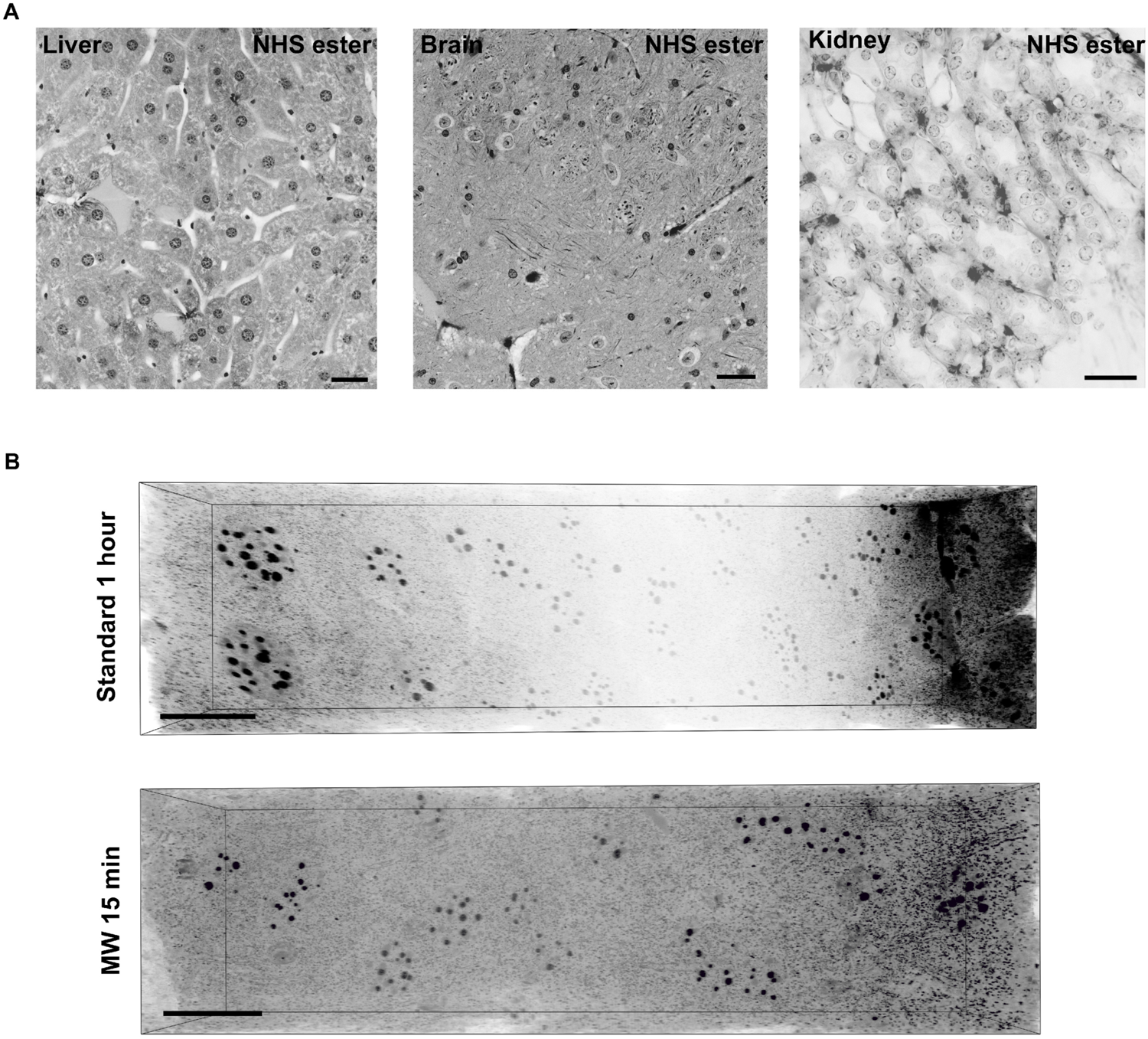
BOOST morphological staining strategy for tissue sections. **(A)** Tile scans of tissue sections (*i.e.*, liver, brain kidney) stained with ATTO 488 NHS ester showing homogeneous staining. **(B)** Microwaved-assisted NHS ester staining (EF = 8.9×) improved the staining uniformity compared with a standard 1-hour NHS staining (EF = 8.9×). Scale bar (biological scale): 20 µm (A); 12 μm (B).

**Fig. S5:**
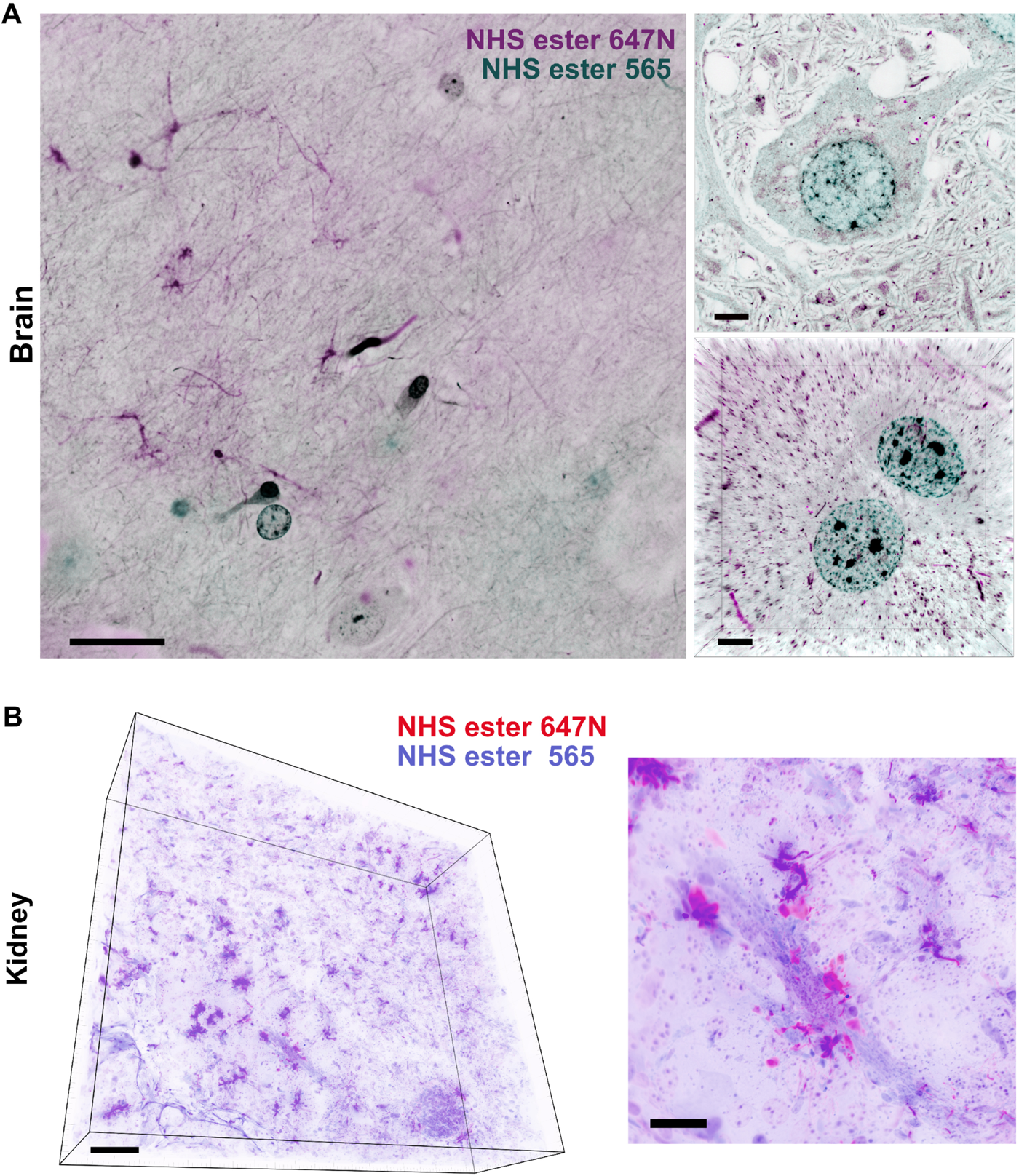
Demonstration of dual proteomic stain strategy for tissue sections. **(A)** A tile scan, zoomed-in region, and a 3D visualization of a brain section stained with dual ATTO 565 NHS ester and ATTO 647N NHS ester showing differential contrast patterns, EF = 9.1×. **(B)** 3D visualization and magnified 3D view of expanded kidney sections stained with ATTO 565 NHS ester and ATTO 647N NHS ester highlighting morphology of different structures, EF = 8.7×. Scale bars (biological scale): 10 µm (A, *left*); 2 µm (A, *right*); 40 µm (B, *left*); 20 µm (B, *right*).

**Fig. S6:**
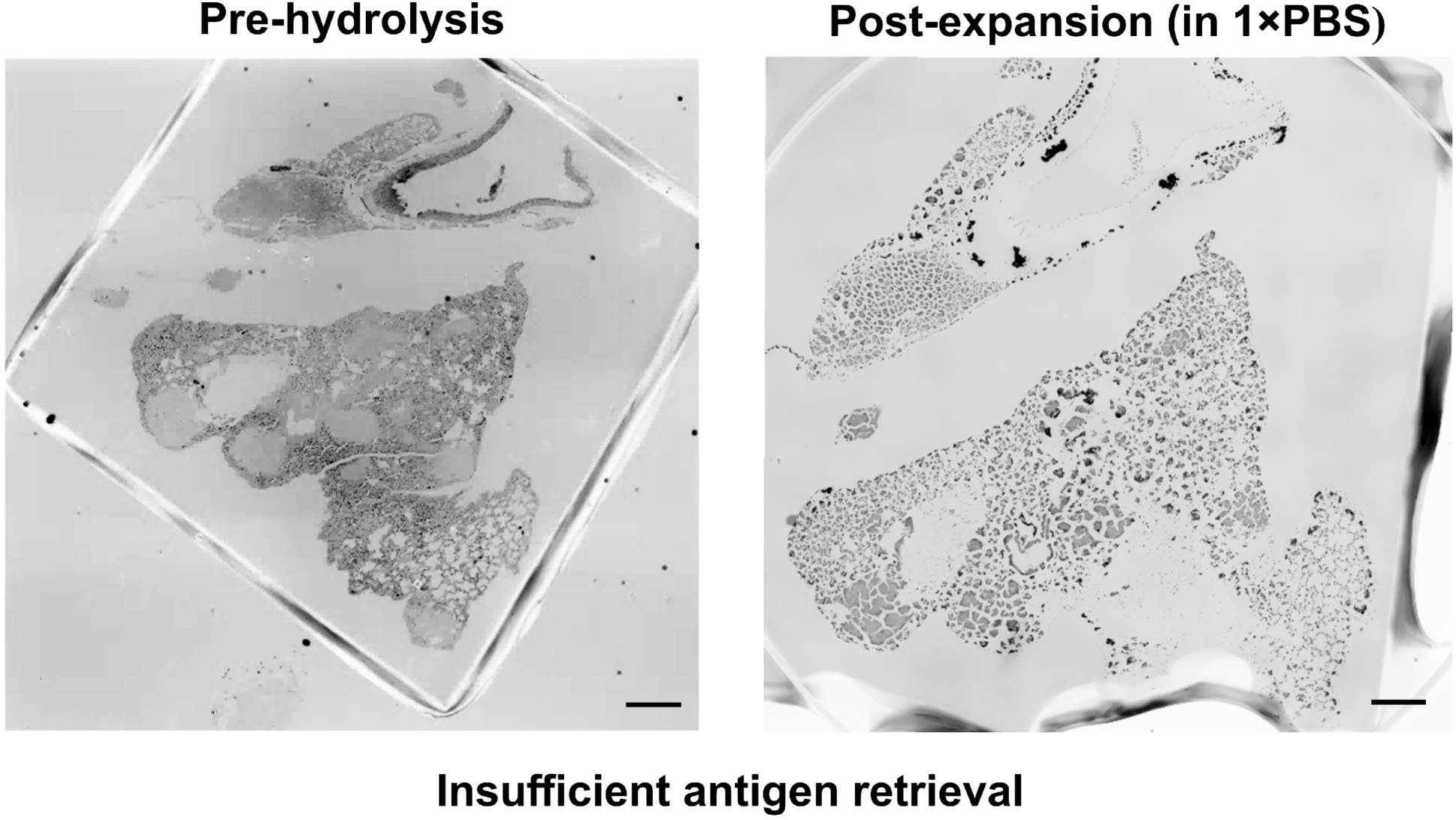
Insufficient antigen retrieval (microwave 5 minutes) resulted in fractured expanded FFPE samples. Images of the same gel-embedded FFPE section before hydrolysis and after hydrolysis in PBS (EF=2.3×). Scale bars (biological scale): 1 mm.

**Fig. S7:**
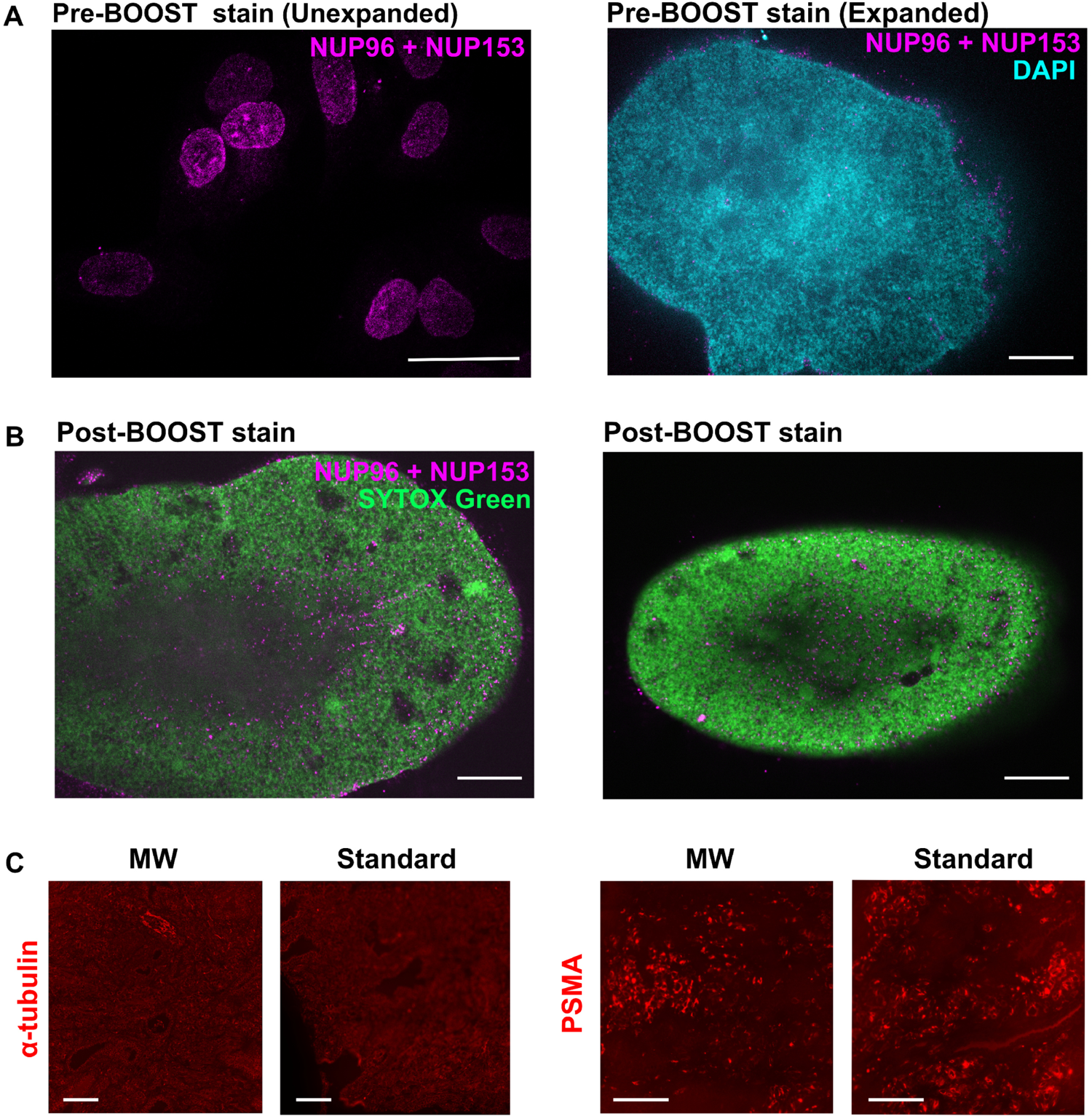
Developing BOOST immunostaining strategy. **(A)** Immunostaining of NPCs with anti-NUP153 and anti-NUP96 antibodies on fixed U2-OS cells (*left*) followed by BOOST processing and DAPI staining (*right*), EF = 9.4×. **(B)** Immunostaining of NPCs of fixed U2-OS cells (*two shown here*) after BOOST denaturation with anti-NUP153 and anti-NUP96 antibodies, EF = 9.6×. The nucleus is stained with SYTOX Green. **(C)** Microwave-assisted immunostaining and standard immunostaining of human prostate cancer FFPE samples after BOOST denaturation generated consistent results. Images were taken in PBS, EF = 2.5×. Scale bars (biological scale): 20 μm (A, *left*); 2 μm (A, *right*); 2 μm (B); 200 μm (C).

**Fig. S8:**
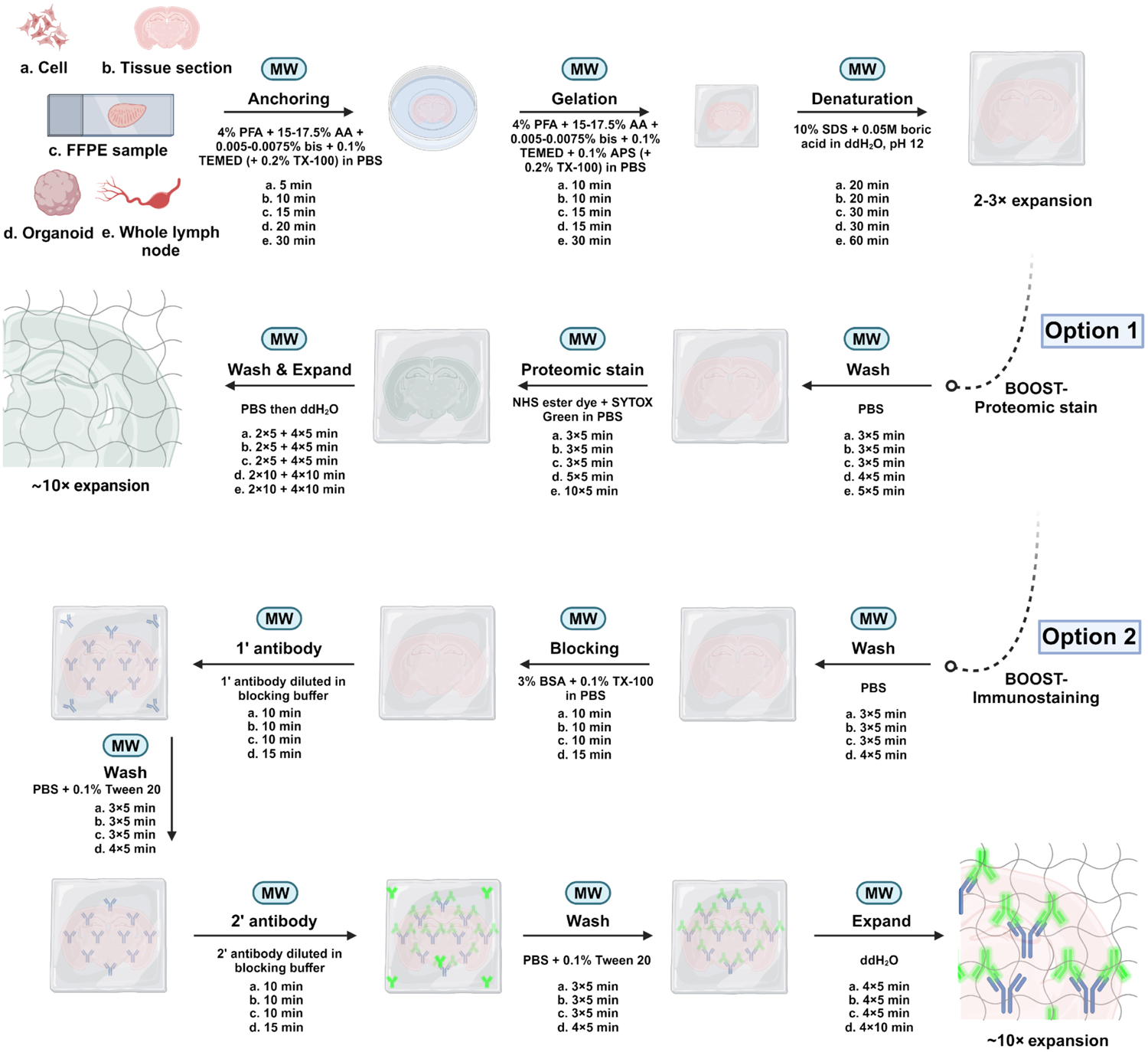
Summary of the BOOST workflows. BOOST workflow on five sample types (*i.e.*, cells, tissue sections, FFPE sample, large 3D organoids, intact lymph node) with the reagents and processing times listed. Two staining options are tested, including microwave-assisted morphological staining with NHS esters with DNA dyes and microwave-assisted immunostaining for visualizing proteins of interest.

**Fig. S9:**
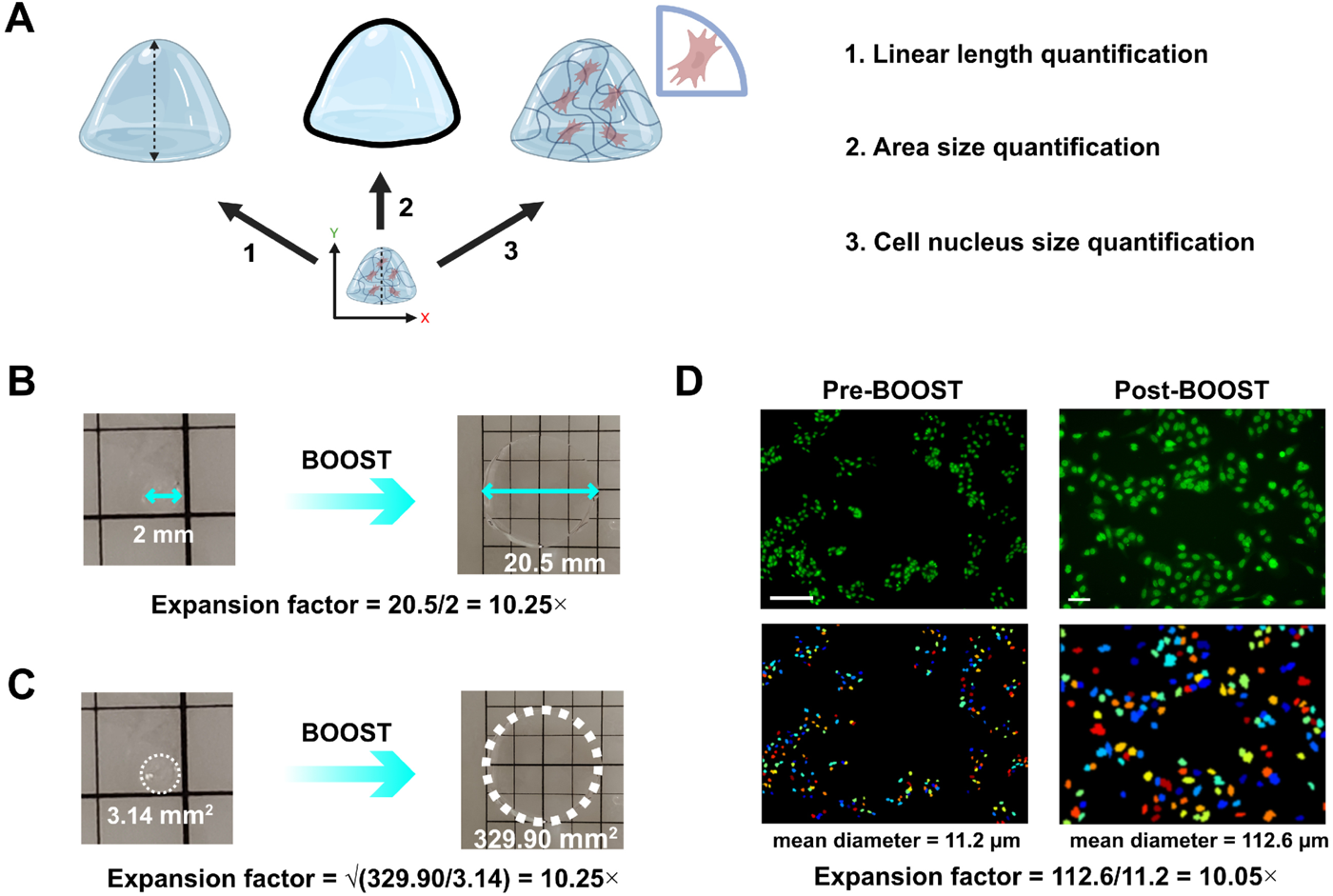
Methods to determine the EF. **(A)** Three strategies are used in this manuscript to determine the EF. **(B)** Expansion factor determination by measuring the expansion of linear length (usually the longest) before and after BOOST. **(C)** EF determination is done by measuring the gel (for cell samples) or specimens (for tissue section, FFPE sample, and large specimens) area before and after BOOST. **(D)** EFdetermination by measuring the sizes of the stained nucleus (n > 50) for each sample before and after BOOST treatment. Scale bar (biological scale): 100 µm (D, *left*); 20 µm (D, *right*).

## Methods

### Chemicals

acrylamide (A9099, Sigma-Aldrich), N,N’-Methylenebis(acrylamide) (146072, Sigma-Aldrich), ammonium persulfate (A7460, Sigma-Aldrich), TEMED (1610801, Bio-Rad), sodium chloride (S3014, Sigma-Aldrich), paraformaldehyde (15700, EMS), glutaraldehyde (16501, EMS), sodium dodecyl sulfate (L3771, Sigma-Aldrich), sodium hydroxide (S5881, Sigma-Aldrich), boric acid (A01298, 3A), ATTO 488, 565, 647N NHS esters (ATTO-TEC), SYTOX Green (S7020, Thermo Fisher), tris base (T6066, Sigma-Aldrich), ethylenediaminetetraacetic acid disodium salt (E5134, Sigma-Aldrich), poly-L-lysine solution (P8920, Sigma-Aldrich), Sigmacote (SL2, Sigma-Aldrich), DAPI (62248, Thermo Fisher), xylene (UN1307, DUKSAN), ethanol absolute (UN1170, Scharlau), Phosphate-buffered saline (PBS) (Gibco 10010023), Bovine serum albumin (BSA) (9998S, ENERGY), Triton X-100 (X100, Sigma-Aldrich), Tween20 (P9416, Sigma-Aldrich), sodium acrylate (408220, Sigma-Aldrich), 4-hydroxy TEMPO (E020145, Energy).

### Antibodies

rabbit polyclonal anti-α-tubulin (11224-1-AP, Proteintech), rabbit polyclonal anti-NUP98-NUP96 (12329-1-AP, Proteintech), rabbit polyclonal anti-NUP153 (14189-1-AP, Proteintech), rabbit polyclonal anti-PSMA/GCPII (13163-1-AP, Proteintech), rabbit monoclonal anti-beta IV tubulin antibody (EPR16776, Abcam), Alexa Fluor 647-conjugated donkey anti-rabbit IgG (ab150063, Abcam), Alexa Fluor 568-conjugated donkey anti-rabbit IgG (ab175470, Abcam),

### Cultured cell samples

U2-OS cells were cultured in DMEM medium supplemented with 10% FBS and 1% penicillin/streptomycin at 37°C in the presence of 5% CO_2_. To prepare cells for fixation, imaging, and expansion, cells were seeded on 12 mm × 12 mm coverslips and cultured until they reached 50-60% confluency for fixation. Cells on coverslips were normally fixed using 4% PFA and 0.1% GA in PBS buffer for 10 minutes. For NUP 98-96 & NUP153 immunostaining, a different fixation strategy, as reported previously, was used to better preserve nuclear pores.^34, 42^ Briefly, cells were firstly prefixed using 2.4% PFA in PBS for 30 seconds, followed by incubation with 0.4% Triton X-100 in PBS for 3 minutes and two additional 5-minute washes in PBS on a rocking platform. Subsequently, the cells were fixed with 2.4% PFA in PBS for 20 minutes at room temperature and then washed twice for 5 minutes with PBS on a rocking platform. A second permeabilization step was carried out using a 0.2% Triton X-100 in PBS for 10 minutes, followed by two additional 5-minute washes in PBS on a rocking platform.

### Mouse tissue section and lymph node samples

Mice were anesthetized using 2% Xylazine : 10% Ketamine : PBS mixed in 1:2:9 and perfused transcardially with 20 mL of PBS and 20 mL of cold fixative solution (4% PFA in PBS). Tissues (*i.e.*, brain, liver, kidney, intact lymph nodes) were then harvested and fixed for 48-72 hours with the same fixative solution at 4°C. Fixed tissues were then sectioned to 100 µm-thick slices using a vibratome (easiSlicer, TED PELLA). The protocol of the mouse experiment was approved by The Committee on the Use of Live Animals on Teaching and Research at the University of Hong Kong.

### Mouse FFPE samples

The lung tumor was established by intravenously injecting 5×10^5^ - 1×10^6^ mouse LLC cells into C57B6/J mice. The lung tissue was collected after 4 weeks of injection. The protocol of the mouse experiment was approved by The Committee on the Use of Live Animals on Teaching and Research at the University of Hong Kong. Tissues were then fixed in 4% PFA in PBS for at least 24 hours. Fixed tissues were then dehydrated and processed as follows: 80% ethanol for 15 minutes, 95% ethanol for 30 minutes for 3 times, 100% ethanol for 45 minutes for 3 times, toluene for 15 minutes for twice, and pre-warmed paraffin for 45 minutes for 3 times. Then samples were embedded in fresh paraffin.

### Human FFPE samples

All prostate cancer paraffin tissues were obtained from the Third Affiliated Hospital of Southern Medical University and approved by the ethics committee of the institution. The pathological diagnosis of the patients was confirmed by two experienced pathologists. Briefly, after surgical resection of the tumor, the samples were fixed with formalin for dehydration immediately, and then embedded with paraffin. 5 μm thickness sections were prepared and mounted on glass slides for subsequent analysis.

### Lung airway organoid samples

The lung organoids were cultured in an expansion medium and passaged every 2 to 3 weeks with a ratio between 1:3 to 1:10, as described previously.^43^ The lung organoids were cultured in the expansion medium for 7-10 days after passaging. To generate apical-out airway organoids, lung organoids were recovered from matrigel and then suspension-cultured in proximal differentiation medium (PneumaCult™-ALI medium + 10 μM Y-27632 + 10 μM DAPT + 100 μg/mL primocin) in Nunclon Sphera culture plate (Thermo Scientific, Waltham, MA, USA) for 14 days. Differentiated airway organoids were then fixed with 4% PFA in PBS for 1 hour at room temperature before processing.

### Microwave set-up

A PELCO BioWave^®^ Pro+ Microwave Processing System (36700, TED PELLA) was used for all the microwave processing in BOOST, with ColdSpot Plus and EM Pro Vacuum Chamber used for specified steps. All the detailed microwave parameters for each step are included in Supplementary Table 1.

### Immunostaining for cultured cells

Cells were washed twice with PBS at room temperature, 5 minutes each, on a rocking platform. Then, samples were permeabilized with PBS containing 0.2% Triton X-100 for 10 minutes at room temperature and washed three times with PBS, 5 minutes each, either on a rocking platform or under microwave irradiation (Supplementary Table 1). Then, samples were blocked using 3% BSA in PBS for 5 minutes under programmed microwave irradiation (Supplementary Table 1), followed by incubation with primary antibodies diluted (1:200 dilution) with 1% BSA in PBS for 5 minutes under programmed microwave irradiation (Supplementary Table 1), then washed three times with PBS, 5 minutes each. Finally, samples were incubated with fluorophore-conjugated secondary antibodies (1:250 dilution) diluted with 1% BSA in PBS for 5 minutes under programmed microwave irradiation (Supplementary Table 1) and washed in PBS for three times, 5 minutes each. Samples will be used for BOOST.

### FFPE sample preparation

FFPE sections mounted on glass slides were deparaffined. Briefly, slides were baked in the oven for 1 h at 60°C, incubated in fresh xylene for 10 minutes twice, and then processed at room temperature in the following flow, 3 minutes for each step: xylene × 3, 100% ethanol × 3, 95% ethanol × 1, 70% ethanol × 1, ddH_2_O × 1, PBS × 3. Antigen retrieval was performed with the slide placed in a microwavable container with the antigen retrieval buffer (Tris/EDTA pH 9) and irradiated with microwave for 30 minutes using the pre-set program (Supplementary Table 1). The samples were cooled to room temperature and then washed with PBS three times.

### Gelation chamber design

#### For sample with thickness < 300 μm (i.e., cells, tissue sections, FFPE, organoids)

Two strips of 300 μm-thick silicone sheet (pre-treated with ethanol to remove impurities) were placed on the parafilm-wrapped or Sigmacote-treated glass slides with appropriate distance left in between for the sample inclusion. The gelation chamber will produce the final gel with a thickness of 300 μm.

#### For thick samples (i.e., organoids, whole lymph nodes)

Silicone sheet of appropriate thickness was selected, or several 300 μm-thick silicone sheets could be stacked for a deeper chamber. Then the sheet or stacked sheets were punched with a hole of appropriate size sufficient to cover the entire sample volume.

### BOOST monomer infiltration, anchoring, and gelation

#### Cultured cells

Cultured cells on the coverslip was placed in a 12-well plate, incubated with anchoring solution (15% w/v AA, 0.005% w/v bis, 0.65M sodium chloride (NaCl), and 4% PFA in PBS) for 5 minutes under programmed microwave irradiation (Supplementary Table 1). Then, the coverslip was placed on the gelation chamber on ice, with the gelation solution (15% w/v AA, 0.005% w/v bis, 0.65M NaCl, 4% PFA, 0.1% w/v TEMED and 0.1% w/v APS in PBS) being mixed well and injected into the space between. The chamber was transferred to the microwave processor for 10 minutes under programmed microwave irradiation (Supplementary Table 1) for gelation. The coverslip with the gel could then be lifted off the chamber, and the gel was cut into a desired size.

#### Tissue sections

The 100-μm vibratome-cut section was first washed in PBS supplemented with 0.2% Triton X-100 three times, 5 minutes each. Then it was placed in anchoring solution (17.5% w/v AA, 0.0075% w/v bis, 0.65M NaCl, 4% PFA in PBS) and incubated for 10 minutes under programmed microwave irradiation in a vacuum environment (Supplementary Table 1). Then, the tissue was transferred to the gelation chamber on ice. A coverslip was placed on the silicone strips to cover the tissue. The gelation solution (17.5% w/v AA, 0.0075% w/v bis, 0.65M NaCl, 4% PFA, 0.1% w/v TEMED and 0.1% w/v APS in PBS) was mixed well and injected into the space between, and transferred to the microwave processor for 10 minutes under programmed microwave irradiation in vacuum environment (Supplementary Table 1) for gelation. The coverslip with the gel could then be lifted off the chamber, and the gel was cut into a desired size.

#### FFPE sections

After FFPE pre-processing step, the slide with FFPE section was cut to a desired size, placed in infiltration/anchoring solution (17.5% w/v AA, 0.0075% w/v bis, 0.65M NaCl, and 4% PFA in PBS) and incubated for 15 minutes under programmed microwave irradiation in a vacuum environment (Supplementary Table 1). Then, the slide was placed on ice, and two 300-μm silicone strips were placed near the tissue section to construct a chamber on the same slide. A Sigmacote-treated coverslip was placed over the tissue section, and the gelation solution (17.5% w/v AA, 0.0075% w/v bis, 0.65M NaCl, 4% PFA, 0.1% w/v TEMED, and 0.1% w/v APS in PBS) was mixed well and injected into the space between. The chamber was then transferred to the microwave processor for 15 minutes under programmed microwave irradiation (Supplementary Table 1) for gelation. The coverslip with the gel could then be lifted off the chamber, and the gel was cut into a desired size.

#### Organoids

The fixed organoids were permeabilized for 5 minutes using PBS + 0.5% Triton X-100. Monomer anchoring was then performed in anchoring solution (15% w/v AA, 0.005% w/v bis, 0.65M NaCl, and 4% PFA in PBS) for 20 minutes under programmed microwave irradiation in a vacuum environment (Supplementary Table 1). Then, the tissue was transferred on the gelation chamber on ice. The gelation solution (17.5% w/v AA, 0.0075% w/v bis, 0.65M NaCl, 4% PFA, 0.1% w/v TEMED, and 0.1% w/v APS in PBS) was mixed well and injected into the chamber space to cover the sample and a coverslip was placed on the silicone sheet to cover the chamber. The chamber was transferred to the microwave processor for 15 minutes under programmed microwave irradiation in vacuum environment (Supplementary Table 1) for gelation. The coverslip with the gel could then be lifted off the chamber, and the gel was cut into a desired size.

#### Whole lymph node

The fixed lymph node was permeabilized for 10 minutes using PBS + 0.5% Triton X-100. Monomer anchoring was then performed in anchoring solution (17.5% w/v AA, 0.0075% w/v bis, 0.65M NaCl, and 4% PFA in PBS) for 30 minutes under programmed microwave irradiation in a vacuum environment (Supplementary Table 1). Then, the tissue was transferred on the gelation chamber on ice. The gelation solution (17.5% w/v AA, 0.0075% w/v bis, 0.65M NaCl, 4% PFA, 0.1% w/v TEMED and 0.1% w/v APS in PBS) was mixed well and injected into the space to cover the sample and a coverslip was placed on the silicone sheet to cover the chamber. The chamber was transferred to the microwave processor for 30 minutes under programmed microwave irradiation in a vacuum environment (Supplementary Table 1) for gelation. The coverslip with the gel could then be lifted off the chamber, and the gel was cut into a desired size.

### BOOST denaturation and hydrolysis conversion

#### Culture cells and tissue sections

The gel was washed with PBS, then transferred with a paintbrush, into a microwavable vessel containing denaturation buffer (10% w/v SDS, 0.05M boric acid in ddH_2_O, pH 12) and processed immediately under microwave irradiation for 20 minutes (Supplementary Table 1). Then, the gel was washed in large volumes of PBS supplemented with 0.1% Tween 20 (PBST) for at least twice, either on a rocking platform or under microwave irradiation (Supplementary Table 1), 5 minutes each.

#### FFPE sections

The slide with the gel was washed with PBS, then placed into a microwavable vessel containing denaturation buffer (10% w/v SDS, 0.05M boric acid in ddH_2_O, pH 12) and processed with programmed microwave irradiation for 30 minutes (Supplementary Table 1). The gel would normally detach from the slide in the first 5 minutes, the slide could be removed, and the gel was hydrolyzed for the rest of 25 minutes. Then, the gel was washed in large volumes of PBST for at least twice, 5 minutes each.

#### Organoids/ Whole lymph nodes

The gel with the organoid/lymph node embedded was washed with PBS, then placed into a microwavable vessel containing denaturation buffer (10% w/v SDS, 0.05M boric acid in ddH_2_O, pH 12) and processed with programmed microwave irradiation for 30/60 minutes (Supplementary Table 1). Then, the gel was washed in large volumes of PBST for at least twice, 10 minutes each.

### BOOST post-denaturation NHS ester staining

#### Cultured cells, tissue sections, and FFPE sections

The denatured gel, after thorough washing, was incubated with ATTO NHS esters diluted in 1:200 in PBS and/or SYTOX Green dyes diluted in 1:500 for 15 minutes under microwave irradiation (Supplementary Table 1). Then, the gel was washed in large volumes of PBST for at least twice either on a rocking platform or under microwave irradiation (Supplementary Table 1), 5 minutes each.

#### Organoids/ Whole lymph nodes

The denatured gel after through washing, was incubated with ATTO NHS esters diluted in 1:200 in PBS and/or SYTOX Green dyes diluted in 1:500 for 25-50 minutes under microwave irradiation (Supplementary Table 1). Then, the gel was washed in large volumes of PBST for at least twice, 10 minutes each.

### BOOST immunostaining after hydrolysis and denaturation

#### Cultured cells, tissue sections, FFPE sections and organoids

The denatured gel after thorough washing, was incubated in the blocking buffer (3% BSA with 0.2% Triton X-100 in PBS) under microwave irradiation for 10-15 minutes (Supplementary Table 1). Then, the gel was incubated with primary antibodies diluted (1:100 dilution) with 1% BSA + 0.1% Triton X-100 in PBS for 10 minutes under programmed microwave irradiation (Supplementary Table 1). The gel was then washed three times with excessive PBST, 5 minutes each, either on a rocking platform or under microwave irradiation (Supplementary Table 1). The gel was finally incubated with fluorophore-conjugated secondary antibodies (1:150 dilution) in 1% BSA + 0.1% Triton X-100 in PBS for 10 minutes under programmed microwave irradiation (Supplementary Table 1). The gel was then washed for at least three times with excessive PBST, 5 minutes each.

### BOOST expansion

For cultured cells, tissue sections, FFPE sections, organoids and whole lymph nodes, large volume of ddH_2_O was exchanged every 5-10 minutes, either on a rocking platform or under microwave irradiation (Supplementary Table 1) until fully expanded, usually four exchanges would be sufficient.

### Gel mounting and imaging

The bottom of plate/dish used for imaging was coated with poly-L-lysine to prevent gel drifting during imaging. Four microscope systems were employed. Epifluorescence images were captured on Keyence X810, using 4×/0.13 NA air (Fig. 2G, 2H, S6), 20×/0.45 NA air (Fig. S7C) and 60×/1.40 NA oil objectives (Fig. 3D). Confocal microscopy and structured illumination microscopy images were captured either using a ZEISS LSM900 upright microscope equipped with Zeiss Plan-Neofluar 2.5×/0.085 NA air (Fig. 4B, 4E), Zeiss N-Achroplan 10×/0.3 NA water immersion (Fig. 4B, 4C, 4F), Zeiss Plan-Apochromat 40×/1.0 NA water immersion (Fig. 2E, 2F, 3C, 4C, S1C, S2A, S2B, S3A, S4A, S5A) objectives or a Vt-iSIM inverted microscope equipped with Olympus UPLSAPO60XW 60×/1.20 NA water immersion objective (Fig. 2C, 2D, 2G, 2H, 3E, 3G, S3B, S6, S7A, S7B). In addition, tested tile images were captured using a LiTone LBS Light-sheet Microscope equipped with a Nikon 16×/0.80 NA water dipping objective (Fig. S5B).

### Assessing hydrolysis conditions in protein and morphology preservation

The workflows were adapted as previously reported^25^ except for the hydrolysis step modified into three conditions (*i.e.,* pH 10, pH 11, and pH 12) for the evaluation of morphological difference and protein retention under 20-minute microwave. Fixed liver slices were first incubated in a monomer solution (30% w/v AA, 0.01% w/v bis, 0.65 M NaCl, 4% w/v PFA, and 0.1% w/v TEMED in PBS) at room temperature for 3 hours. Then, the tissue was transferred to the gelation chamber (monomer solution with 0.1% w/v APS). A coverslip was placed on the silicone strips to cover the tissue and the fresh gelation solution was injected into the space between. The chamber was then placed at room temperature for 40 minutes to complete gelation, and the gel was cut into appropriate sizes and recovered in PBS. For the modified hydrolysis, hydrolysis buffer (0.2M SDS, 0.05M boric acid in DI water) for three pH conditions: pH 10, pH 11, and pH 12, was prepared separately. The recovered gel was then transferred with a paintbrush, into a microwavable vessel containing denaturation buffer of specified pH value and processed immediately under microwave irradiation for 20 minutes (Supplementary Table 1). The denatured gel was then washed in PBS four times, 1 hour each. The washed gel was stained with ATTO 565 NHS ester diluted 1:200 in PBS for 1 hour at room temperature, then washed in ddH_2_O three times, 15 minutes each, followed by an exchange of ddH_2_O every hour to a fully expanded state for imaging.

### Optimization of microwave-assisted anchoring

The workflows were adapted from previously published ^2^ except for the anchoring step modified into four conditions (*i.e.*, no anchoring, 10-minute microwave, 20-minute microwave, and the standard TREx protocol of 1-hour incubation) for the evaluation of AcX effectiveness under microwave. For standard TREx anchoring, liver sections were incubated in 100 μg/mL AcX diluted in PBS for 1 hour at room temperature. For modified microwave-assisted anchoring, liver sections were incubated in 100 μg/mL AcX diluted in PBS and processed in a microwave processor for 10 minutes and 20 minutes (Supplementary Table 1), respectively. After the anchoring step, the sections were rinsed in PBS. Later incubated in TREx gelation solution (10.34% w/v SA, 14.22% w/v AA, 0.005% w/v bis, 0.15% w/v APS, 0.15% w/v TEMED, 0.0015% w/v TEMPO in PBS) for 20 minutes on ice. Then, the tissue section was transferred on the gelation chamber on ice. A coverslip was placed on the silicone strips to cover the tissue. The fresh gelation solution was then injected into the space between. The chamber was placed in a 37°C incubator for 1 hour to complete gelation, and the gel was cut into appropriate sizes and recovered in PBS. Gel was then placed into TREx disruption buffer (5% w/v SDS, 0.2M NaCl, 0.05M Tris in ddH_2_O, pH 7.5) in a tube and the tube was transferred to an incubator set at 80°C for 3 hours. The denatured gel was then recovered in 0.4M NaCl followed by two washes in PBS, 30 minutes each. The washed gel was stained with ATTO 565 NHS ester diluted 1:200 in PBS, then washed in ddH_2_O three times, 15 minutes each, followed by an exchange of ddH_2_O every hour to a fully expanded state for imaging.

### Standard post-denaturation immunostaining for FFPE sample

The denatured FFPE gel after thorough washing, was incubated in the blocking buffer (3% BSA with 0.2% Triton X-100 in PBS) at room temperature for 3 hours. Then, the gel was incubated with primary antibodies diluted (1:100 dilution) with 1% BSA + 0.1% Triton X-100 in PBS overnight at 4°C. The gel was then washed three times with excessive PBST, 30 minutes each, on a rocking platform. The gel was finally incubated with fluorophore-conjugated secondary antibodies (1:150 dilution) in 1% BSA + 0.1% Triton X-100 in PBS for 3 hours at room temperature. The gel was then washed for at least three times with excessive PBST, 30 minutes each.

### Expansion factor determination

We used three strategies to determine expansion factors (Fig. S9). First being the linear measurement of the longest distance of the unexpanded gel and expanded gel, second being the square root of the area ratio of the gel size in two states and lastly being the cell nucleus size quantification based on pre-BOOST SYTOX Green, NHS ester staining or NUP immunostaining of unexpanded and expanded images.

### Distortion and RMSE analysis

The unexpanded and expanded image distortion was evaluated using B-spline transformation in Elastix, and RMSE was analyzed according to protocols and script offered by Vaughan’s group.^44^

### Deformation index (DI) analysis

Blank gel was formed in a gelation chamber with a thickness of 800 μm, a 4 mm diameter region was punched to produce a circular gel, then hydrolysis was performed (only for BOOST formulations), and gel was expanded to a fully expanded state. To calculate the DI, the gel was cut in half and stood against the ruler wall (with the curved side facing downwards to allow the flat surface to deform), images were taken and later processed in ImageJ for measurement of the best-fit deformed circle. Two BOOST formulations were used (17.5% w/v AA / 0.0075% w/v bis; 15% w/v AA /0.005% w/v bis), and the original TREx formulation was also evaluated (10.34% w/v SA, 14.22% w/v AA, 0.005% w/v bis, 0.15% w/v APS, 0.15% w/v TEMED in PBS).

---

**Table 1.**
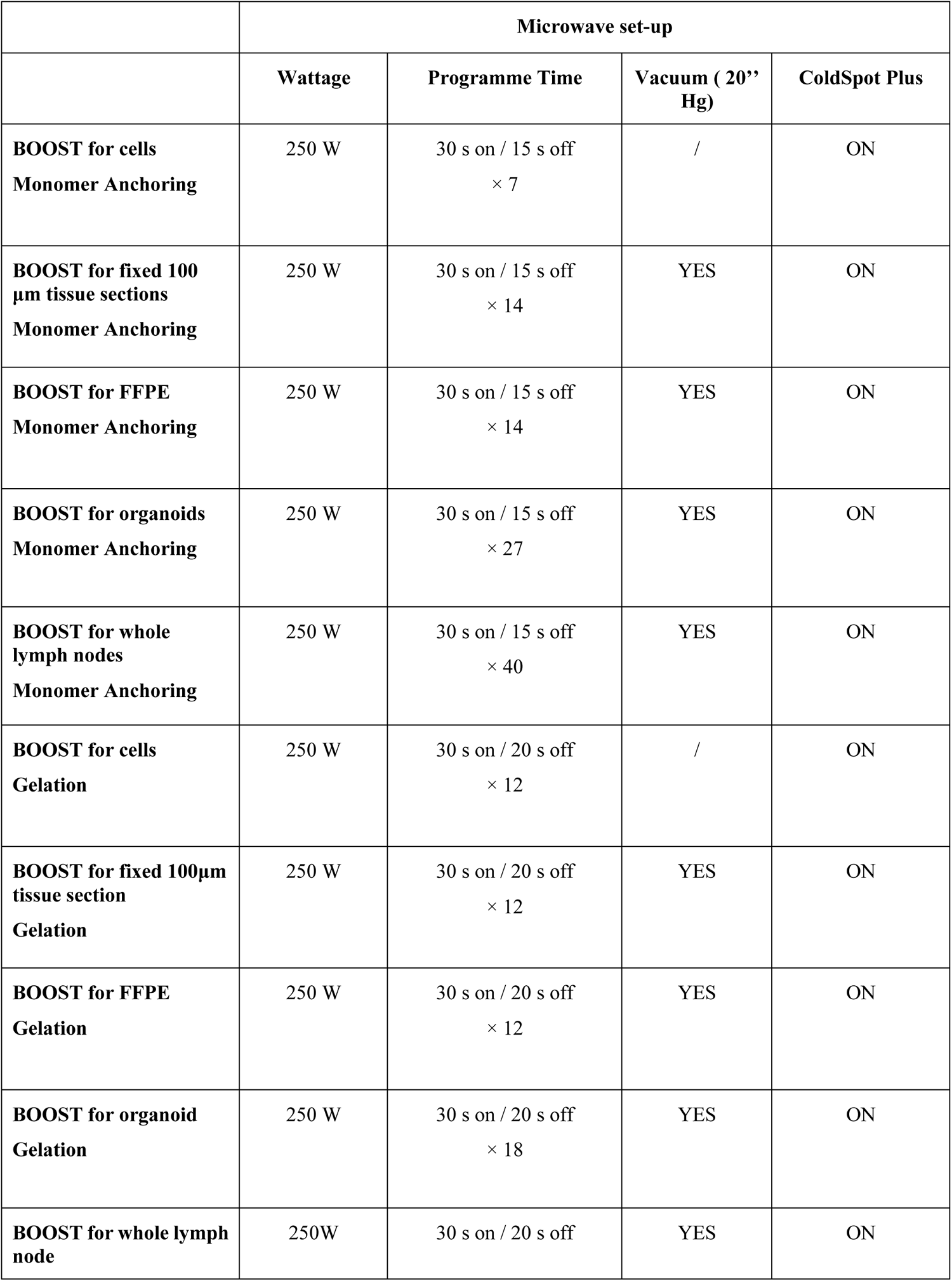
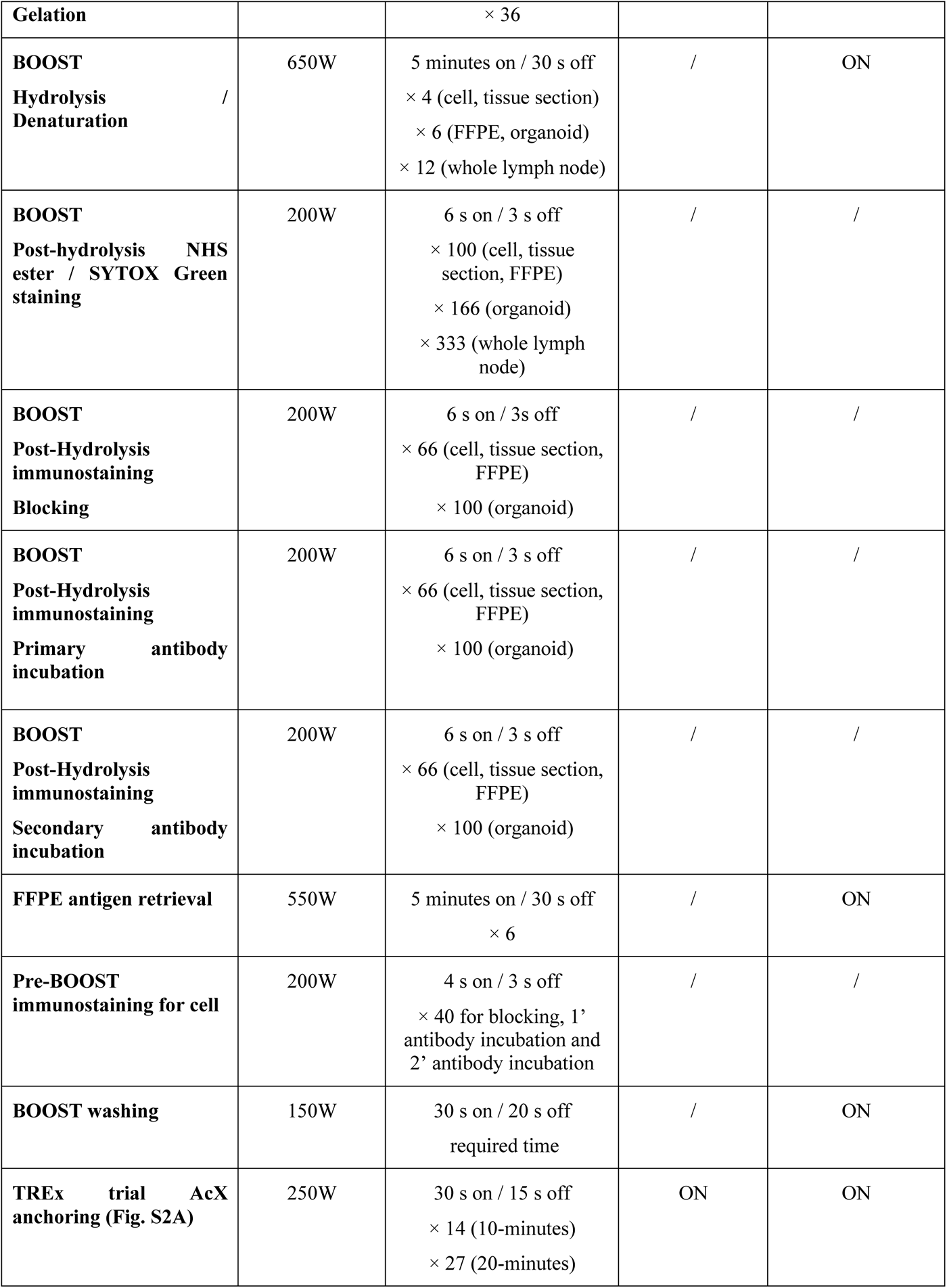
Microwave settings for BOOST processes.

